# Octopi: Open configurable high-throughput imaging platform for infectious disease diagnosis in the field

**DOI:** 10.1101/684423

**Authors:** Hongquan Li, Hazel Soto-Montoya, Maxime Voisin, Lucas Fuentes Valenzuela, Manu Prakash

## Abstract

Access to quantitative, robust, yet affordable diagnostic tools is necessary to reduce global infectious disease burden. Manual microscopy has served as a bedrock for diagnostics with wide adaptability, although at a cost of tedious labor and human errors. Automated robotic microscopes are poised to enable a new era of smart field microscopy but current platforms remain cost prohibitive and largely inflexible, especially for resource poor and field settings. Here we present *Octopi*, a low-cost ($250-$500) and reconfigurable autonomous microscopy platform capable of automated slide scanning and correlated bright-field and fluorescence imaging. Being highly modular, it also provides a framework for new disease-specific modules to be developed. We demonstrate the power of the platform by applying it to automated detection of malaria parasites in blood smears. Specifically, we discovered a spectral shift on the order of 10 nm for DAPI-stained Plasmodium falciparum malaria parasites. This shift allowed us to detect the parasites with a low magnification (equivalent to 10x) large field of view (2.56 mm^2^) module. Combined with automated slide scanning, real time computer vision and machine learning-based classification, Octopi is able to screen more than 1.5 million red blood cells per minute for parasitemia quantification, with estimated diagnostic sensitivity and specificity exceeding 90% at parasitemia of 50/ul and 100% for parasitemia higher than 150/l. With different modules, we further showed imaging of tissue slice and sputum sample on the platform. With roughly two orders of magnitude in cost reduction, Octopi opens up the possibility of a large robotic microscope network for improved disease diagnosis while providing an avenue for collective efforts for development of modular instruments.

**One sentence summary:** We developed a low-cost ($250-$500) automated imaging platform that can quantify malaria parasitemia by scanning 1.5 million red blood cells per minute.

## Introduction

Lack of cost-effective diagnostics is a major hurdle in global fight against infectious disease, specially in resource poor settings [1]. This leaves our world in a highly vulnerable position with therapeutic drugs being either overused, leading to drug resistant strains or not accessible to people who actually need these treatments [2]. Since health care is delivered around the world in a tiered structure, local context such as high cost, lack of trained personal or low throughput of many available diagnostics tests plays a large detrimental role on quality of delivered health-care [3].

Because of the versatility and wide adoption of manual microscopy [4] and its role in direct visual identification of parasites [5], it remains a WHO gold standard for numerous diseases [1]. Despite technological advancements in related fields, the practice of conventional manual microscopy has remained largely unchanged over the last half century and suffers from several drawbacks. With an average lab technician spending 6 to 8 hours imaging and examining slides per day, human fatigue has been identified as a crucial factor in reduced efficiency in microscopy based diagnostics [6]. With heavy disease burden, number of patient samples that need to be processed, even at small primary health centers, can often supersede the capacity of laboratory workers [7]. The above listed limitations for microscopy are not fundamental, and can be circumvented with field implementation of low-cost, motorized microscopes combined with computer-based automated detection.

Low-cost field microscopy has made tremendous strides in the last decade [8], both towards access and implementing application-specific capabilities [9, 10, 11, 12, 13, 14]. New microscopy techniques such as Fourier ptychographic microscopy [15] and lens-free on-chip microscopy [16, 17, 18] have also been developed to tackle some of the hurdles of conventional microscopy in diagnostics settings. These platforms and techniques have demonstrated a wide range of applications [19, 20] but high throughput diagnosis of malaria has remained out of reach.

Despite all the resources invested, malaria remains to be a highly deadly disease. In the year of 2017, there were 219 million cases and nearly 435,000 deaths, majority of them occurring due to *Plasmodium falciparum*, a strain of malaria widely spread across the world [21]. Two most widely used diagnostic tests are antigen-based Rapid Diagnostic Test (RDT) and microscopic examinations of blood smears. In the same year, 276 million RDTs were sold and more than 208 million patients were tested by microscopy, whereas estimated needs for testing was well over 1 billion [22]. While RDT is easy to use, it cannot quantify parasitemia, stays positive post treatment for up to a month [23] and can create false negative due to HRP2/3 gene deletions [24, 25]. Manual microscopy, on the other hand, is labor intensive and in practice the performance is often compromised. Commercial slide scanning and detection systems show promise [26, 27] but are currently expensive. With persisting high burden of malaria [28] and the rise of drug resistance strains [29, 30, 31], affordable, high-throughput and quantitative diagnostic tests are urgently needed.

Here we present *Octopi*, a low-cost ($250-$500), portable (below 3 kg), reconfigurable and automated imaging platform for disease diagnosis in resource constrained settings. To enable versatility of the platform and its adoption for different diseases, we take a highly modular approach where the platform can be configured with different disease-specific modules. On this platform, we demonstrate automated slide scanning with multimodal imaging with two imaging heads that support a range of magnifications.

In particular, we report a spectral shift on the order of 10 nm for DAPI-labeled *P. falciparum* malaria parasites when compared to often confounding DAPI-labeled platelets in patient samples. This discovery enables us to integrate three channels of information (bright-field, fluorescence and spectral) for automated detection of *P. falciparum* parasites with a low magnification imaging module. Large field of view afforded by this module, combined with automated slide scanning and image processing, allows screening of more than 1.5 million red blood cells per minute for infections, which is 120 times faster compared to traditional manual microscopy [32]. We further implement a machine learning classifier and obtain anticipated performance of higher than 90% specificity and sensitivity for parasitemia of 50 parasites per *µl* and 100% sensitivity and specificity for parasitemia of 150 parasites per *µl*. Our results suggest that low-cost automated multimodal microscopy combined with machine learning tools have the potential to address the unmet needs for diagnosis of malaria and many other diseases.

## Results

### Automated imaging platform with modular design

The imaging platform consists of completely separable modules that fall into 5 categories: imaging, slide scanning, transillumination, oblique angle laser illumination and control & computation (Fig. 1A). When setting up the imaging platform (Fig. 1B), preassembled modules snap to each other due to embedded magnets (see mounting of the imaging head in Movie S1 for example). Since screws are not necessary for the connections, the imaging platform can be rapidly reconfigured.

**Fig. 1.**
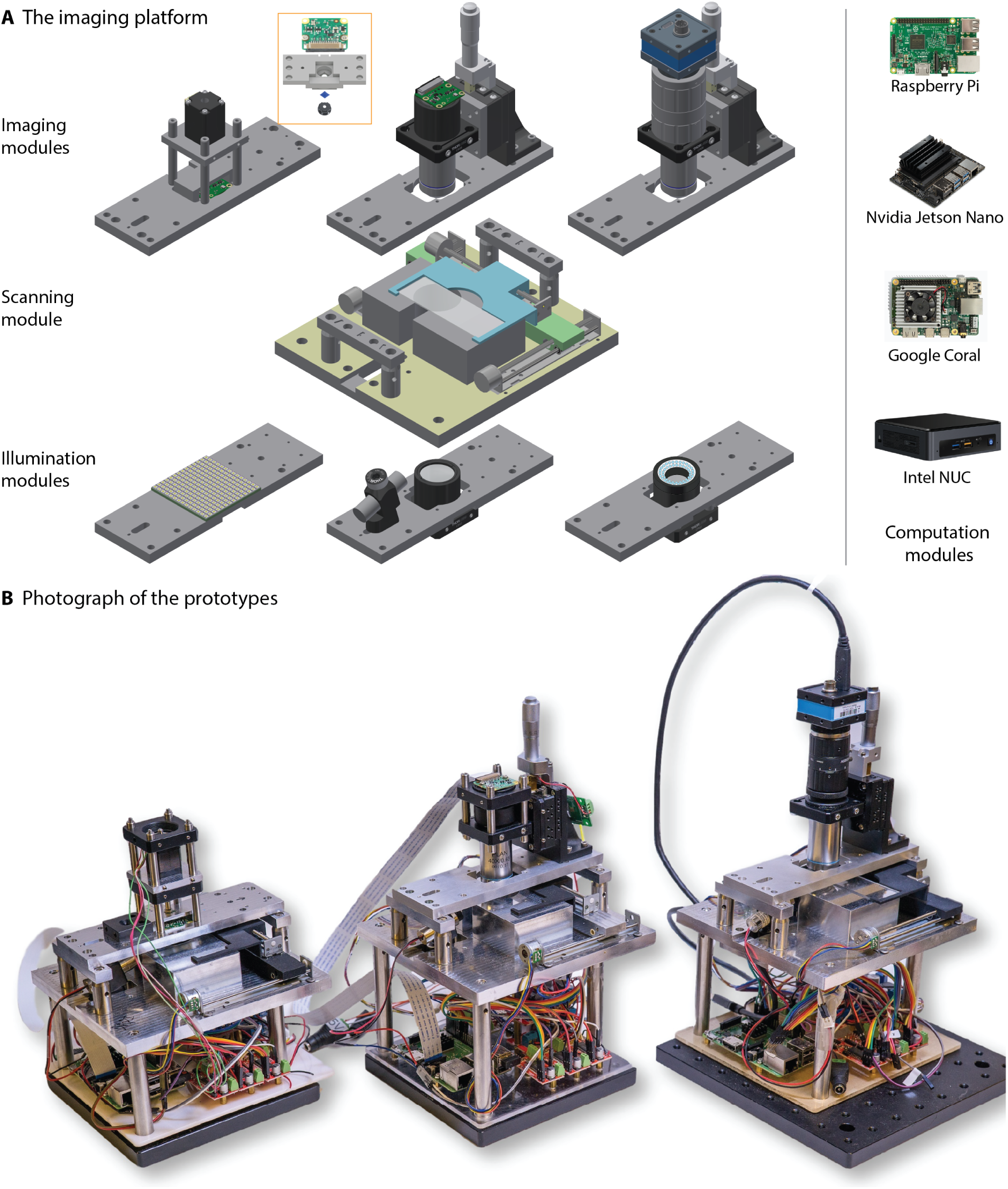
Reconfigurable high-throughput imaging platform. **(A)** Construction of the modular imaging platform. The left column shows three different imaging modules (top row), a motorized scanning module, and three different illumination modules (bottom row). In the *low mag imaging module* (top left), a captive linear actuator is used for focus actuation. In the *high mag imaging module* (top middle and top right), piezoelectric stacks combined with micrometers are used for focus actuation, where the micrometer can be replaced with a captive linear actuator to motorize coarse adjustment. Inset shows the construction of the low-mag imaging module sub-assembly, which consists of a pi-camera, a long pass interference filter and another cellphone lens. For different applications, sub-assemblies with different configurations should be switched as a whole, in contrast to the *high mag imaging module*, where objectives, filters, tube lens and cameras can be individually switched. The right column shows some examples of currently available portable computing devices that can be used as the computation module. **(B)** A photograph showing three *Octopi* prototypes with different imaging modules optimized for different applications.

We designed the platform with a combination of standard and custom parts, with choices being made to optimize performance, size, cost, and ease in prototyping and iterative development. For example, we made the imaging and transillumination module compatible with the standard cage and lens tube system, which allowed us to quickly implement different configurations. For the custom parts that form the backbone of the microscope, we chose CNC machining with 6061 aluminum over other manufacturing options for the rigidity of metal, the tight tolerance of the machining process and the low surface roughness of the finished parts. CNC machining also has favorable cost-volume scaling: at the manufacturing quantity of 10, the price is already comparable with 3D printing.

To facilitate wide adoption of the imaging platform, including in resource limited settings, cost is imposed as an important design constraint during development. Through careful choices of parts and their arrangements, we were able to keep the starting unit cost of the imaging platform to about $700 for volume of 10 units. Without significantly altering the design, the cost reduces to $350 for volume of 100 units and $250 for volume of 1000 units (table S1).

#### The imaging modules

We implemented two different imaging modules, one with low magnification (*low mag imaging module* and one with high magnification (*high mag imaging module*). The *low mag imaging module* is based on the reversed lens configuration, where two multi-element cell-phone lenses are used as objective and tube lens in the infinity-corrected configuration [33]. To enable fluorescence imaging, an interference long pass filter diced into the size of 3 mm x 3 mm can be placed in between the two lenses (Fig. 1A inset). The CMOS sensor (Pi Camera based on Sony IMX219), lenses and optional filter are assembled around a CNC machined part as a permanent assembly. Because the cost of the parts is low, for different filters or lens combinations, different permanent assembly can be made. This eliminates the needs for users to handle small and intricate parts and helps keep the optical train free from dusts and contamination. The f-number of the lenses used in our implementation is 2.0, which translates to numerical aperture of 0.25, typical of 10X objectives. With condenser-based transillumination for bright field and oblique angle laser illumination for fluorescence, we got Nyquist-limited resolution of 2.5 *µm* (2.3 times the object side pixel size) over field of view of more than 1.6mm x 1.6mm (Fig. 2A). By using different pairs of lenses, diffraction limited resolution (0.92 *µm*) of a 10X/0.3NA objective lens may be achieved.

**Fig. 2.**
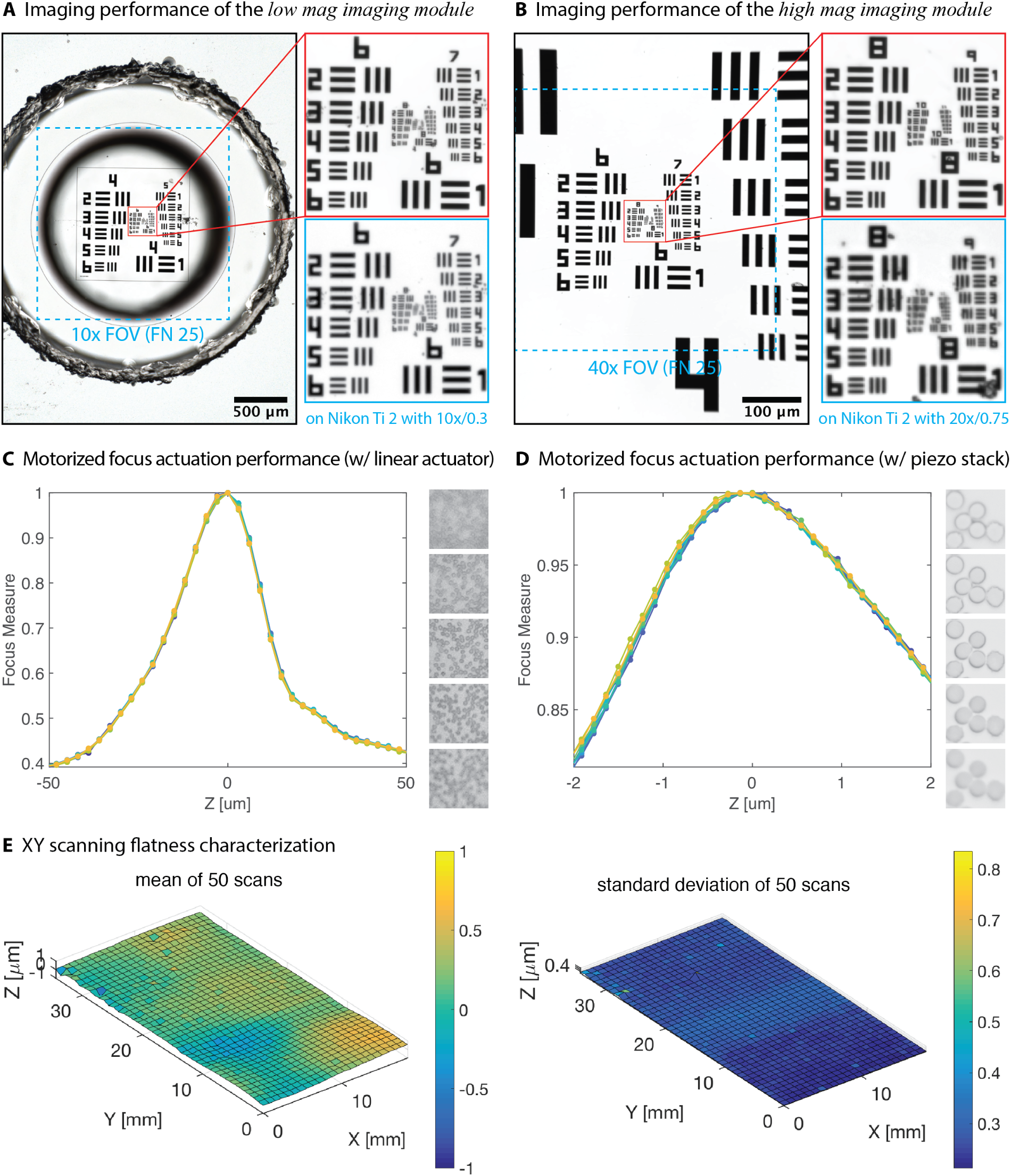
Characterization of the imaging platform. **(A-B)** Images of a 1951 USAF resolution obtained on *Octopi* with the *low mag imaging module* and the *high mag imaging module* and their comparisons with images obtained on a Nikon Ti2 microscope. The image obtained with the *high mag imaging module* (configured with 40x/0.65 objective) is better resolved that its counterpart obtained on the Nikon Ti2 with an aprochromatic 20x/0.75 objective and an additional 1.5x magnification because of its smaller object side pixel size (0.202 um compared to 0.367 um). Images were denoised by a pretrained FFDNet denosiser [93], see Fig. S13 for images before denoising. **(C)** Motorized focus actuation performance of the *low mag imaging module* (using captive linear actuators) and the *high mag imaging module* (using piezoelectric stacks). Plotted are focus curves (focus measure vs commanded z position) for 10 repeated z-stacks. Step size of 3 um and 137 nm is used for the captive linear actuator and the piezoelectric stack respectively. The high degree of overlap between the curves suggest reliable and repeatable focus actuation. Example images are 12 *µ*m and 1.1 *µ*m apart in z. **(D)** Characterization of XY scanning flatness for scanning module using an ultra-flat glass slide. Mean and standard deviation of measured top surface z-positions for 50 XY scans are plotted. The overall standard deviation is below 400 nm and is limited by the measurement setup, suggesting excellent stage flatness. Similar result was obtained with a normal microscope glass slide and plotted in Fig. S3.

The *high mag imaging module* makes use of standard infinity-corrected microscope objectives. Depending on required sensitivity and frame rate, Pi Camera or standard industrial camera may be used, with M12 lens or C-mount imaging lens acting as tube lens. Notably, with starting price below $100, industrial cameras with low light CMOS sensors can offer peak quantum efficiency of more than 70% and readout noise as low as 1.1 e- (see Fig. S1), rivaling the performance of high-end scientific cameras. Using Pi Camera, a f = 25mm M12 lens and a 40x/0.65 Plan Achromatic objective, with the same illumination used for the low mag module, Nyquist-limited resolution of 0.46 *µm* can be achieved with field of view of 0.4mm x 0.4mm (Fig. 2B).

High-throughput automated imaging requires robust auto-focus. In the *low mag imaging module* we implemented motorized focusing with a captive linear actuator. The step size of the linear actuator is 1.5 *µm*, which is sufficiently small compared to the depth of focus of more than 8 *µm*. Motorized focus adjustment for the *high mag imaging module* has more stringent requirements, given the depth of focus is as small as 1 *µm* for a 40x/0.65 objective. As a solution, we combined a low-cost piezo stack actuator and a standard linear translation stage with extended contact ball bearings/crossed roller bearings. The piezo stack actuator used has travel of 11.2 *µm* and step size of 2.73 *nm* when used with a 12-bit digital to analog converter (see movie S2 for demo of focus actuation with this implementation). To test the performance of the motorized focus actuation for the low and high mag module, we acquired series of z-stacks of blood smears and plotted the computed focus measures vs the commanded z-position (Fig. 2 C,D). That the curves lie on top of each other indicates excellent reliability and repeatability. Utilizing the dependence of focus measure on z position, we implemented contrast-based auto-focus. Alternatively, with small modifications in illumination, different single-shot focus-finding approaches [34, 35] can be used for faster focus.

#### Illumination modules

The bright field transillumination module consists of a LED panel, a diffuser and an NA = 0.79 condenser. The diffuser is placed at the focal plane of the condenser to make the illumination Khler-like. Dark-field illumination for low magnification can be provided simply by a ring of LED, while an LED matrix can be used for quantitative phase imaging, fourier ptychography [15, 36] and single-shot auto-focus [34, 35]. For fluorescence excitation, we make use of oblique angle laser illumination [9]. Used in a wide range of consumer electronics such as Blu-Ray/DVD/CD players, projectors and laser pointers, direct diode laser and diode-pumped solid state lasers that can provide tens to hundreds of mWs optical power are available at a wide range of wavelengths at very low cost. Currently, available wavelengths include 405 nm, 450 nm, 465 nm, 505 nm, 520 nm, 532 nm, 635nm, 650 nm, 780 nm, 808 nm, 1064 nm. Because of the monochromatic nature of the laser, no excitation filter is needed. The use of oblique angle illumination also eliminates the need for dichroic beam splitter, reducing both the overall size and the cost of the setup. In addition, multiple lasers can be used and electronically switched for multiplexing.

#### The scanning module

Motorized slide scanning is crucial for high throughput imaging. Commercial motorized stage for microscopes offers incremental motion as low as a few nanometers but typically cost thousands of US dollars. Realizing that for wide field imaging such level of positioning performance is not needed, we developed a low cost scanning module using lead screw linear actuator that costs as little as $5 per unit. A important performance criterion is the scan flatness, which is the relative z-displacement of the slide at the center of the microscope field of view as the slide is being scanned. Good flatness reduces the need for frequent auto-focus. To ensure good scanning flatness, in our present configuration, the slide rests directly on a CNC machined block and is moved by a slide scanner driven by the lead screw linear actuators. To characterize our stage flatness, we used an ultra-flat quartz coated glass slide (and in another measurement, a normal microscope glass slide) as target and measured with a non-contact sensor the displacement of its top surface as it is being scanned (Fig. S2). The result (Fig. 2E, Fig. S3), which is limited by measurement setup, suggests overall flatness below 400 nm over tens of millimeters.

#### The control & computation module

Raspberry Pi, a single board computer priced at $35, provides a cost-effective way to control the microscope. The linux operating system also makes it easy to take advantage of open source software packages and simplifies development. In the Raspberry Pi-based implementation, the camera is interfaced using the industry standard MIPI camera serial interface, whereas other components are controlled through driver boards and MOSFET switches.

With increasing demands for artificial intelligence at the edge, various low-cost and energy-efficient ASIC chips and embedded systems with optimized hardware for computer vision and machine learning applications have recently emerged. For applications requiring more compute power and/or higher imaging throughput, these platforms can be adopted. In particular, we have implemented and tested image processing and spot detection pipelines on Jetson Nano, a $99 drop-in replacement for Raspberry Pi with 128 CUDA cores. This implementation reduces processing time by more than 50 times and allows processing to be done in real time as slides are being scanned (Fig. S4). Further more, when Windows-only software needs to be used, or more compute power is required, laptops or desktop workstations can also be used.

#### Power consumption

When Raspberry Pi or Jetson Nano are used as the control & computation module, the entire system can be powered from 5V DC power supplies. Either a wall plug AC adapter or a battery pack may be used. For a battery pack with capacity of 20,000 mAh, a single charge can power the microscope for more than 8 hours of continuous use (assuming full power consumption).

### Automated blood smear examination

Blood smear examination is commonly used for diagnosing blood-borne diseases and often requires imaging many microscopic field of views. *Octopi* as a high-throughput imaging platform is particularly suited for these applications. To use the platform, stained blood smear is prepared following the same protocol used for manual microscopy (Fig. 3A). The slide is then loaded to the imaging platform and scanned automatically. For samples stained with fluorescent dyes, in addition to bright field image, a fluorescent image is also taken for each field of view, during which the LED is switched off and the laser is switched on by the controller (movie S1).

**Fig. 3.**
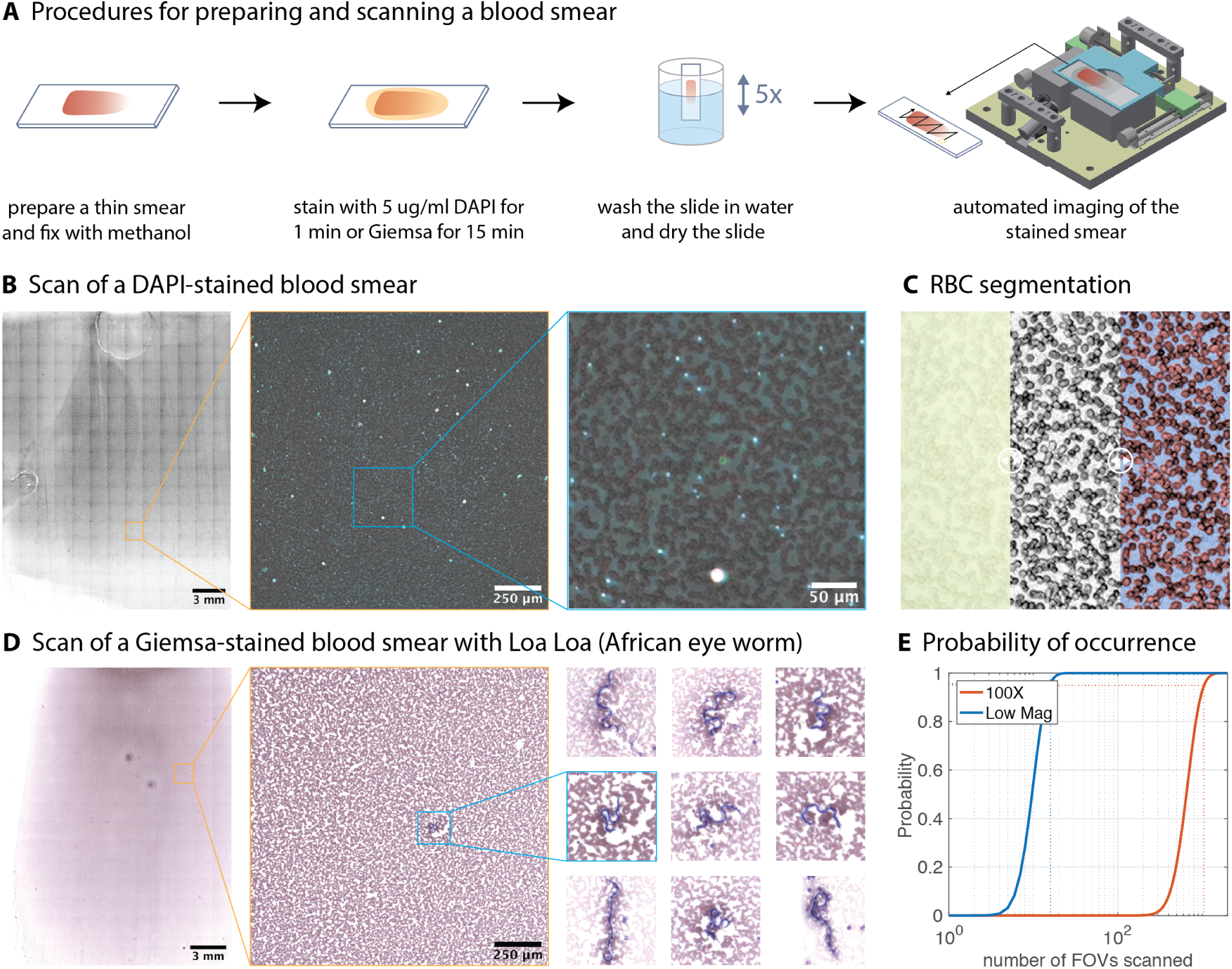
Blood smear examination. **(A)** Procedure for preparing and scanning a blood smear. **(B)** Scan of a DAPI-stained blood smear. From left to right: stitched bright field images, a single FOV with overlaid bright field and fluorescent images, zoomed-in overlay image with arrows pointing to (from top to bottom) a platelet, a reticulocyte and a white blood cell. The smear is made from 4 *µl* of blood and the region being imaged is of size 20.8 mm x 27.2 mm (221 individual field of views), covering more than 90% of the smear. With Raspberry Pi being used as the control & computation module, The scan took 19 minutes. This includes auto-focus using 20-plane z-stacks at the beginning and in the middle of each row that accounts for about 1/3 of the total scan time. When implemented with a control & computation module that has higher bandwidth (such as Jetson Nano), shortening of the total acquisition time to below 4 minutes can be achieved. Besides, in practice, digitizing a much smaller area of the blood smear is often sufficient. **(C)** Illustration of steps for segmentation of red blood cells. Left to right: unprocessed portion, portion with preprocessing applied (illumination correction and contrast adjustment), portion with segmentation masked generated from a neural network overlaid. **(D)** Scan of a Giemsa-stained blood smear with Loa Loa (African eye worm). The 9 zoomed-in images are of size 188 *µ*m *×* 188 *µ*m **(E)** Assuming parasitemia of 100 parasites/ul of blood, probability of more than 10 parasites present vs total number of microscopic fields of view examined. The two curves are for the *low mag imaging module* (field of view: 1.6 mm x 1.6 mm) and for a 100x objective commonly used for malaria diagnosis (field of view: 0.22 mm in diameter). In the calculation, red blood cells is assumed to fill 75% of the field of views. For the probablity to be greater than 95%, on average more than 1058 of the 100x field of views need to be examined, whereas only 16 low mag fields of view is sufficient.

Because of the absence of nucleus in red blood cells, fluorescent dyes that bind to the nucleic acid may be used for staining platelets, white blood cells and many parasites in blood smears with improved contrast compared to bright field stains. The dark field nature of fluorescence imaging also makes it possible to localize stained objects that are below the diffraction limit of the objective being used. This allows us to use low magnification optics that have lower resolutions but larger field view, resulting in higher imaging throughput. Furthermore, in fluorescent imaging, since angular distribution of emitted light is independent of illumination, full numerical aperture of the objective is automatically utilized.

Among many fluorescent nucleic acid dyes, 4’,6-diamidino-2-phenylindole (DAPI) has several attractive properties, including 20-fold fluorescence enhancement upon binding to the AT-region of dsDNA, low-cost (staining a blood smear costs less than $0.02 without reusing the staining solution), and good temperature stability. According to published data sheet [37], DAPI solutions are stable at room temperature for 1 to 2 weeks [38] and at +4*^◦^*C for up to 6 month. In practice, we got similar staining results with DAPI solution (at staining concentration of 5 *µ*g/ml) left in the dark at room temperature for several months. To demonstrate the use of DAPI with the *low mag imaging module* on our platform, we scanned a smear of whole blood that is stained in DAPI solution for one minute. In the resulting images, not only white blood cells, but also reticulocytes and platelets can be easily resolved (Fig. 3B).

To be able to robustly extract fluorescent spots for quantification, we developed a two-step processing pipeline (Fig. S5). The first step uses top-hat transformation to remove background. The second step uses a blob detector based on the Laplacian of Gaussian (LoG) [39, 40] to detect fluorescent spots of different sizes and intensities. Since filtering operations involved in both steps are computationally expensive for CPUs, we implemented a version of the pipeline that takes advantage of CUDA cores in GPU. When deployed on Jetson Nano for detecting platelets (and later, also malaria parasites), we’re able to get per image processing time of around 300 ms, which is more than 50 times faster compared to using a Raspberry Pi 3B+ and 5 times faster compared to using desktop computers (Fig. S4). With scanning speed of 1 field of view per second, this allows equivalent blood smear examination throughput of 3,000,000 red blood cells (or 0.6 *µl* blood) per minute (assuming red blood cells cover 75% area of the field of view).

Directly counting red blood cells can be beneficial for quantitative analysis of the blood and for determining parasitemia in the case of infection, especially when the precise volume of blood being smeared is not known. However, at low magnification, when cells are only stained with fluorescent dyes, segmentation becomes challenging. In particular, because each red blood cell has only 5-7 pixels in diameter and the contrast is not uniform across the cell, simple thresholding or edge detection-based methods do not work well. Hough transforms used for detecting circular objects requires the image to be scaled up, which leads to significant processing overhead, and has trouble detecting red blood cells that have distorted shapes. To address this challenge, we trained a 91-layer fully convolutional DenseNet [41], which gives good performance (Fig. 3C, Fig. S6). By compressing the model through pruning, quantization and other optimizations [42] and deploying it on Jetson Nano, real-time performance can be expected. To further improve throughput, more lightweight model [43, 44] may be trained and ASIC chips may be used [45].

To demonstrate use case for detecting larger parasites using only bright-field imaging, we digitized a Giemsa-stained blood smear with Loa-Loa parasites (Fig. 3(D)). Because the parasite can be identified unambiguously, blood volume limited-detection limit of 0.2-0.5 parasites/*µl* can be achieved. This detection threshold is well below the Sever Adverse Effect (SAE) threshold of 30 parasites/*µl*, above which mass drugs should not be administrated to the individual patient [12]. For screening this particular parasite, video microscopy [12] has also been used and can be configured on *Octopi* to further improve throughput.

The advantage of automated scanning becomes evident by examining the probability of occurrence of a given number of parasites in scanned fields of view. For a hypothetical parasitemia of 100 parasites/*µl*, we plot the probability of more than 10 parasites being present as a function of number of fields of views scanned for two different magnifications (Fig. 3E). We can observe that if enough fields of views are examined, the probability goes to one. Furthermore, in applications where the use of fluorescent dyes and/or pathogen-specific probes renders the morphological features of the detection targets unimportant, or reduces the requirement of optical resolution (such as in detecting DAPI-stained platelets), low magnification can be used in place of high magnification to significantly boost throughput. In the example above, to have at least 10 parasites with more than 95% probability, on average only 16 low mag fields of view are needed, as compared to 1058 fields of views in the case of 100 x oil objective. This calculation assumes that all the targets are in the same plane. For targets that are distributed in 3D, such as in sputum sample or in tissue slices, the increase in throughput is even more significant, given the depth of focus of 55 um, 8.8um, 1.3 um and 0.53 um for 4x/0.1, 10x/0.25, 40x/0.65 and 100x/1.25 oil objectives.

### Automated detection of malaria parasites in thin blood smears

Fluorescence microscopy has been used for sensitive detection of malaria parasites [46, 47, 48]. However the prospect of detection in fixed blood smears at low magnification is hindered by the presence of brightly stained platelets, which are highly abundant (there are typically 250,000 platelets per *µl* blood) and appear similarly in size and brightness as malaria parasites. Yet, *P. falciparum* malaria parasites, which have a 48-hour asexual life cycle, contain not only DNA but also large amount of RNA. This provides an opportunity for differential detection. Previously, it has been shown that the emitted fluorescence red-shifts in DAPI-RNA complexes compared to DAPI-DNA complexes [49], which means that depending on the DNA-RNA ratio cells, overall shift up to about 40 nm can be expected. In fact, this property has been used in enumerating reticulocytes in rodent malaria models [50].

To support the feasibility of differentiating malaria parasites from platelets based on DNA/RNA ratio and its associated spectral shift, we imaged smears of blood from healthy individuals and patients diagnosed with malaria with laser scanning confocal microscopy where spectrum at each pixel is recorded. The results revealed a spectral red shift on the order of 10 nm for ring-stage *P. falciparum* parasites. For better visualization of the results, we mapped the obtained 32-channel spectral stacks to pseudo color images (Fig. 4A, Fig. S7-S8), where the color is determined by centroids of the spectrum, with purple being 485 nm or below and yellow being 510 nm or above. Using the same color code, we plotted the spectrum of selected spots (Fig. S8) in Fig. 4(B), where three clusters emerge. Examining the spots (Fig. 4(C)) we can conclude that the first purple/dark blue cluster (centroid below 495 nm) corresponds to platelets, and that the second green colored cluster (centroid at 495-500 nm) belongs to ring-stage malaria parasites. Because of the absence of distinctive morphological features, the identity of the third cluster where the “yellow” spectrum originate (centroid above 505nm) remain to be determined. Likely candidates for the brighter “yellow” spots include merozoites and trophozoites stages of the *P. falciparum* parasites, as these stages can be stained intensively with RNA-selective dyes[51]. As they’re not observed in uninfected blood, dimmer “yellow” spots can be accounted for by parasites-derived extracellular vesicles, which have been reported to contain RNA and DNA[52, 53, 54, 55].

**Fig. 4.**
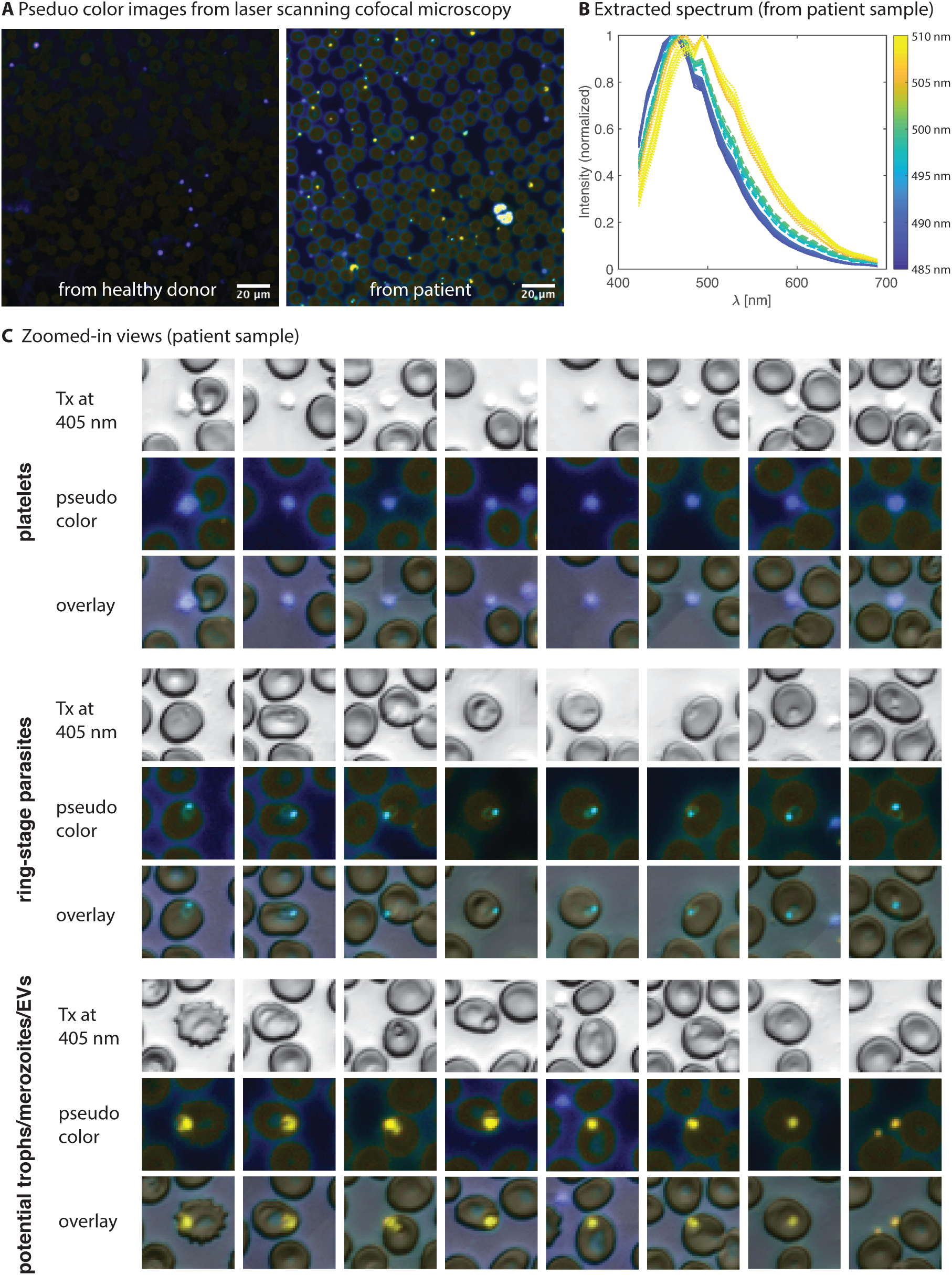
Spectral shift in DAPI-stained *P. falciparum* parasites. **(A)** Pseuduo color images of DAPI stained blood smear from a healthy donor and from a patient diagnosed with malaria. The images are acquired on a Zeiss LSM780 laser scanning confocal microscope with a 20x/0.8 objective and 32 spectral channels. For each pixel, the color is determined according to the centroid of the extracted 32-point spectrum for that pixel **(B)** Extracted spectrum of selected fluorescent spots from image of the patient sample where each spectrum is color-coded according to its spectral centroid in the same way as in (A). **(C)** Zoomed-in views of platelets, ring-stage *P. falciparum* malaria parasites and potential *P. falciparum* malaria trophozoites, merozoites and parasites-derived extracellular vesicles. Images are of size 17 *µ*m *×* 17 *µ*m.

Traditionally, fluorescence microscopy is done with monochrome cameras and band pass filter with relatively narrow pass band for better sensitivity and background suppression. However, in doing so, spectral multiplexing will involve use of multiple filters or point spread function engineering, which adds to the complexity of the system. Here by utilizing a long pass filter and a color CMOS sensors where color filter arrays in the Bayer arrangements are directly integrated on top of the pixels, we are able to obtain spectral information in a single shot. To quantify the performance of this setup, we simulated the spots with spectrum from the average of DAPI-stained platelets and DAPI-stained ring-stage parasites (Fig. 5A, Fig. 5B). In the simulation, spots were assumed to have a Gaussian profile, and both finite pixel size and photon shot noise were taken into account. To get a lower-dimensional representation, the spots are then projected to normalized color space G/B vs R/B, where R/B is the ratio of total red pixel intensity and total blue pixel intensity, and similarly G/B is the ratio of total green pixel intensity and total blue pixel intensity (Fig. 5C). Intriguingly, for spot sizes and signal levels easily achievable, spectral shift as low as 8 nm can result in good separation in the G/B vs R/B space.

**Fig. 5.**
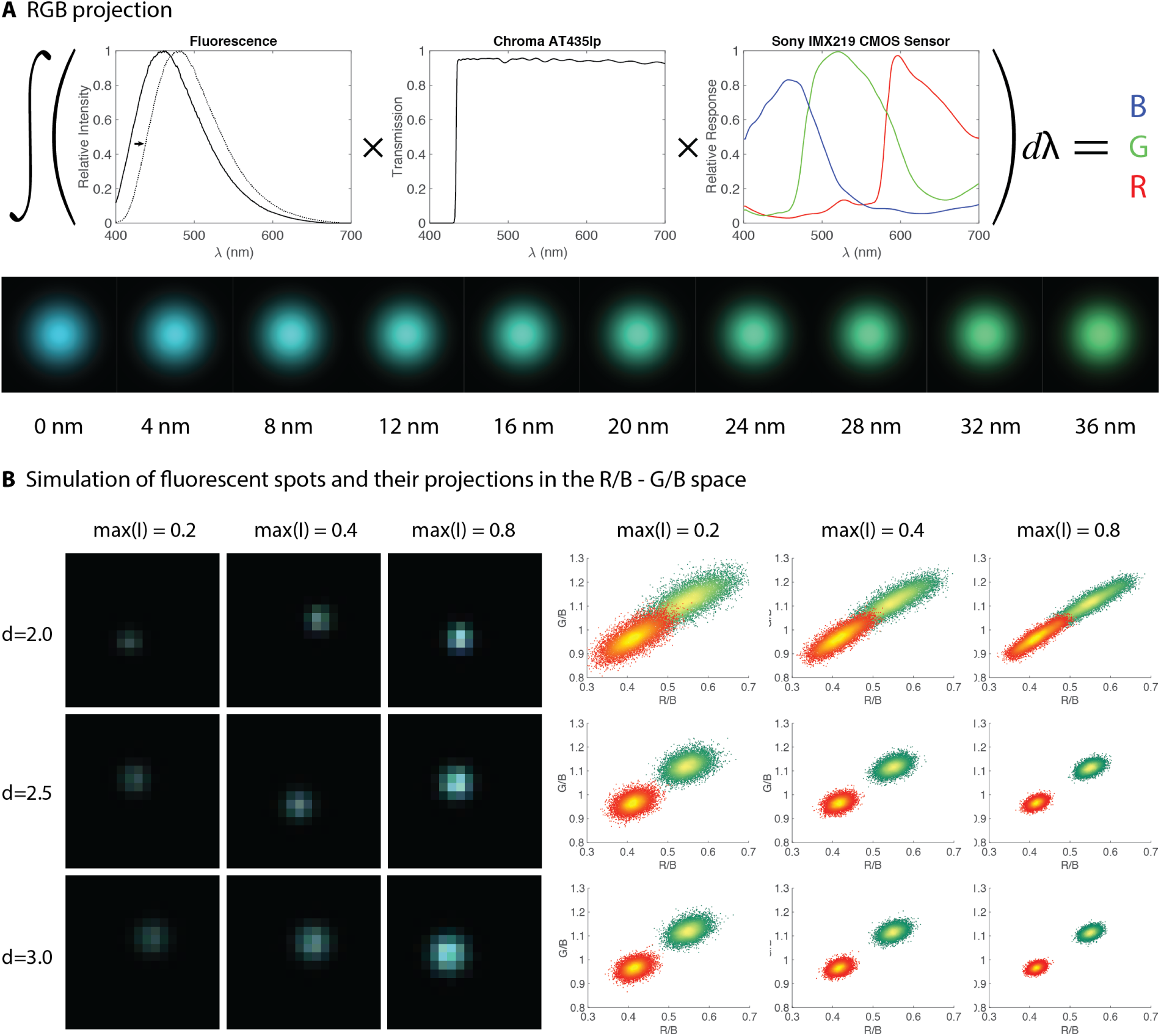
Simulation of three channel spectral imaging with an RGB CMOS sensor. **(A)** The process of converting spectrums to RGB values (top) and the resulting colors for different amount of shift. The original spectrum without shift is the emission spectrum from DAPI. **(B)** Example demosaiced images of simulated fluorescent spots of different diameter and signal to noise level (left) and their projections in the R/B - G/B space (right; the two clusters have spectral separation of 8 nm). For each parameter combination, 10,000 spots are randomly generated for each class. The spots are assumed to have Gaussian profiles and the diameters are their RMS width. The signal level is the expected value of the maximum pixel intensity of the spot. The number is normalized to have max value of 1, which corresponds to the full well capacity of the CMOS sensor. In determining shot noise for the pixel values, peak quantum efficiency conversion gain of the pi camera module is used (peak QE: 70%, conversion gain: 0.2 e-/ADU). The shot noise is modeled by a Poisson process. The position of spots are also randomized. For each spot, R/B is the ratio of total red pixel intensity and total blue pixel intensity, and similarly G/B is the ratio of total green pixel intensity and total blue pixel intensity. The R, G, B pixel values are directly taken from the simulated raw Bayer data.

To show that our imaging platform configured with the *low mag imaging module* has enough sensitivity for detecting DAPI-stained ring-stage parasites, we imaged the same smear of *P. falciparum* culture on *Octopi* and on a high end research microscope (Nikon Ti2 with Prime 95B sCMOS sensor), and one-to-one correspondence of fluorescent spots can be observed (Fig. 6A). Fig. 6B compares a typical overlaid bright-field and fluorescent field of view of *P. falciparum* culture with that of uninfected whole blood, and the color difference between parasites and platelets can be appreciated. To quantify how well parasites and platelets may be told apart, we stained and imaged 8 smears of *P. falciparum* culture and 10 smears of uninfected whole blood, where a total number of 109,355 fluorescent spots from the *P. falciparum* culture and 437,944 fluorescent spots from the uninfected whole blood were detected and extracted using the aforementioned processing pipeline. Projection of randomly selected 10,000 spots into the G/B vs R/B space is plotted in Fig. 6C. Good separation in this scatter plot suggests and that color, as a manifestation of spectral shift, is a robust feature for distinguishing parasites from platelets. The results also suggest the absence of confounding objects in uninfected whole blood.

**Fig. 6.**
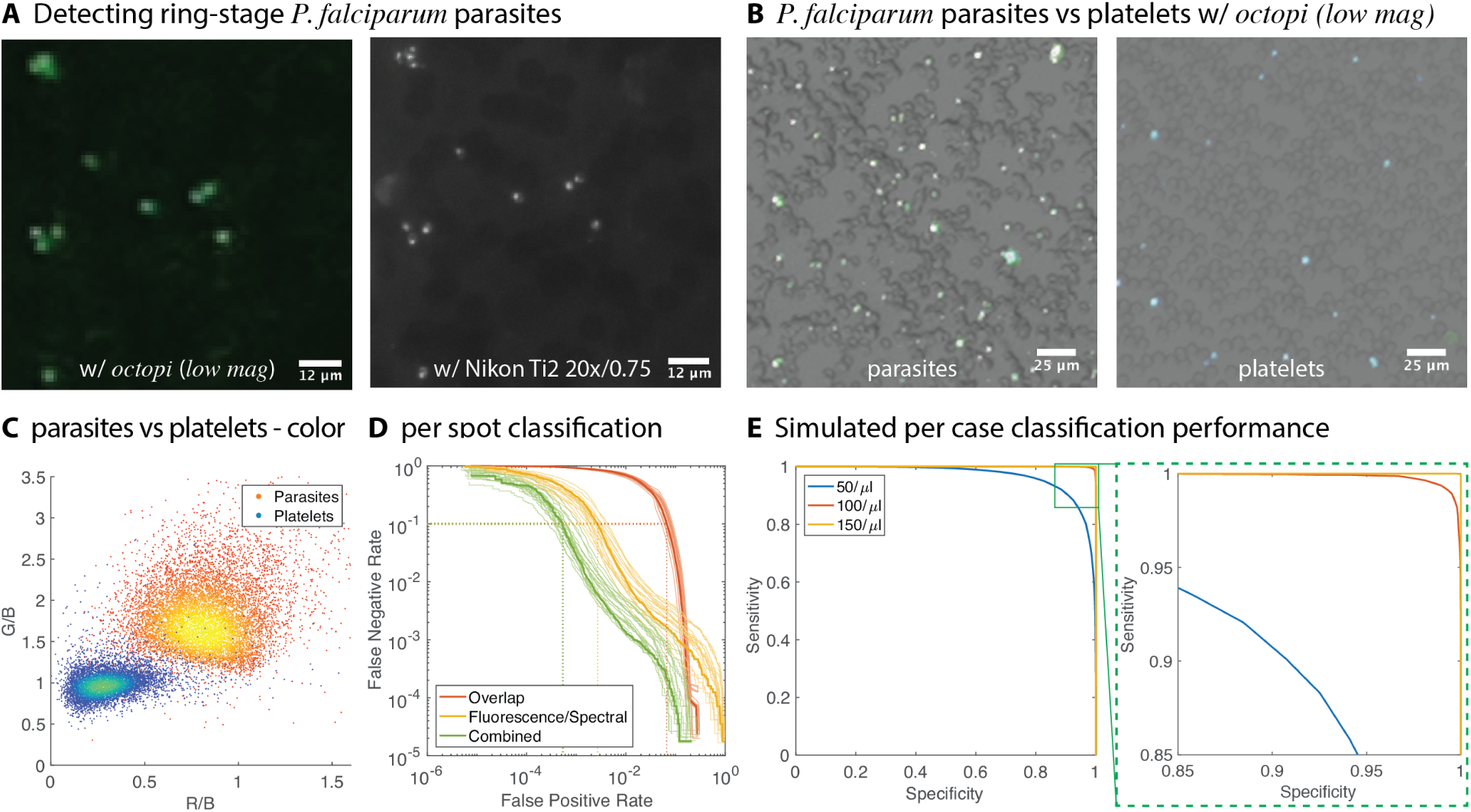
High throughput *P. falciparum* parasites detection at low magnification. **(A)** Image of the same field of view of DAPI-stained smear of *P. falciparum*. culture obtained with the *low mag imaging module* on *Octopi* (left) and on Nikon Ti2, a high-end research grade microscope, with 20x/0.75 apochromatic objective (right). **(B)** Overlaid bright field and fluorescent images of DAPI-stained smears of *P. falciparum*. culture (left) and uninfected whole blood (right) obtained with the *low mag imaging module*. The color difference of fluorescent spots in the two can be observed. We can also observe that the platelets are all outside right blood cells where as most parasites are inside the red blood cells. **(C)** Scatter plot of spots corresponding to parasites and platelets in the G/B vs R/B space. The spots are labeled according to whether they come from the *P. falciparum* culture smears or the uninfected whole blood smears. In the scatter plot, 10,000 randomly chosen spots from each class is shown. **(D)** Per spot classification performance for three different classes of classifier with 20-fold cross validation. The first classifier only uses features extracted from the fluorescent image for classification. The second classifier only uses the amount of overlap between the fluorescent spot and the segmented red blood cells. The third classifier uses both fluorescent features and overlap, which gives 0.05% false positive rate at false negative rate of 10%. **(E)** Simulated per case classification performance assuming per spot FNR of 5 *×* 10*^−^*^4^ and FNR of 10%. 10,000 tests at each parasitemia level were simulated, assuming examination of 0.5 *µ*l blood (2.5 million red blood cells) per test. Each test outputs an estimated parasitemia based on the number of red blood cells scanned and number of parasites detected, and this number is compared with a decision threshold for determining the outcome of the test. For each simulated parasitemia, this decision threshold is varied to obtain the sensitivity vs specificity curve. We note that per case sensitivity and specificity is a measure of performance at low parasitemia. For higher parasitemia (e.g., above 200/*µ*l), estimated parasitemia may be directly used. Compared to RDT, the ability to quantify parasitemia is a strength of microscopy and is useful for evaluating disease severeness and monitoring treatment response.

To automatically detect parasites from extracted spots and obtain diagnostic performance that can be expected with the proposed solution, we built a boosted-tree classifier that takes features from each extracted spots and outputs a class label. The performance of the classifier can be characterized by its False Positive Rate (FPR) and False Negative Rate (FNR), where FPR is the number of platelets misclassified as parasites over the total number of platelets and FNR is the number of parasites misclassified as platelets over the total number of parasites. We found that using combined features from bright-field images and fluorescent images result in the best classification performance (Fig. 6D). Specifically, at FNR of 10%, FPR of 0.05% (average of 20-fold cross validation, range is 0.027%-0.11%, standard deviation is 0.019%) can be achieved. Because both declaration of a smear as negative and quantifying parasitemia in the case of low parasitemia involves scanning a large area and counting a large number of cells, and that brightly-stained platelets are highly abundant, it’s important to choose a decision threshold that gives relatively small per spot FPR. This lowers the chances of misdiagnosing an uninfected case as infected and only has a weak negative influence on sensitivity. With per spot FPR = 5 *×* 10*^−^*^4^ and FNR = 11%, we obtain through Monte Carlo simulations anticipated (per case) sensitivity and specificity of (91%,91%), (99%,99%) and (100%,100%) for parasitemia of 50/ul, 100/ul and 150/ul (Fig. 6E). This simulation assumes platelet count of 250,000/ul, all platelets being brightly labeled and that 0.5 *µl* blood is screened. In the Jetson Nano-based implementation, the time it took from slide being loaded to an answer (including parasitemia, in the case of infection) can be less than 2 minutes.

In certain cases it may be desirable to resolve the morphology of individual parasites. This would further improve sensitivity and specificity, especially for cases with very low parasitemia. This is made possible on our modular platform by using the *high mag imaging module* with a 40x/0.65 objective. We imaged smears of uninfected whole blood (Fig. 7A), lab culture of *P. falciparum* (Fig. 7B) and blood samples from patients diagnosed with malaria (Fig. 7C). The result show that with morphology and/or color, parasites can be easily told apart from platelets. Images of lab *P. falciparum* culture also confirm that many parasites are indeed in their ring-stage, with presence of multiple infections, which is due to the high concentration of parasites in the lab culture.

**Fig. 7.**
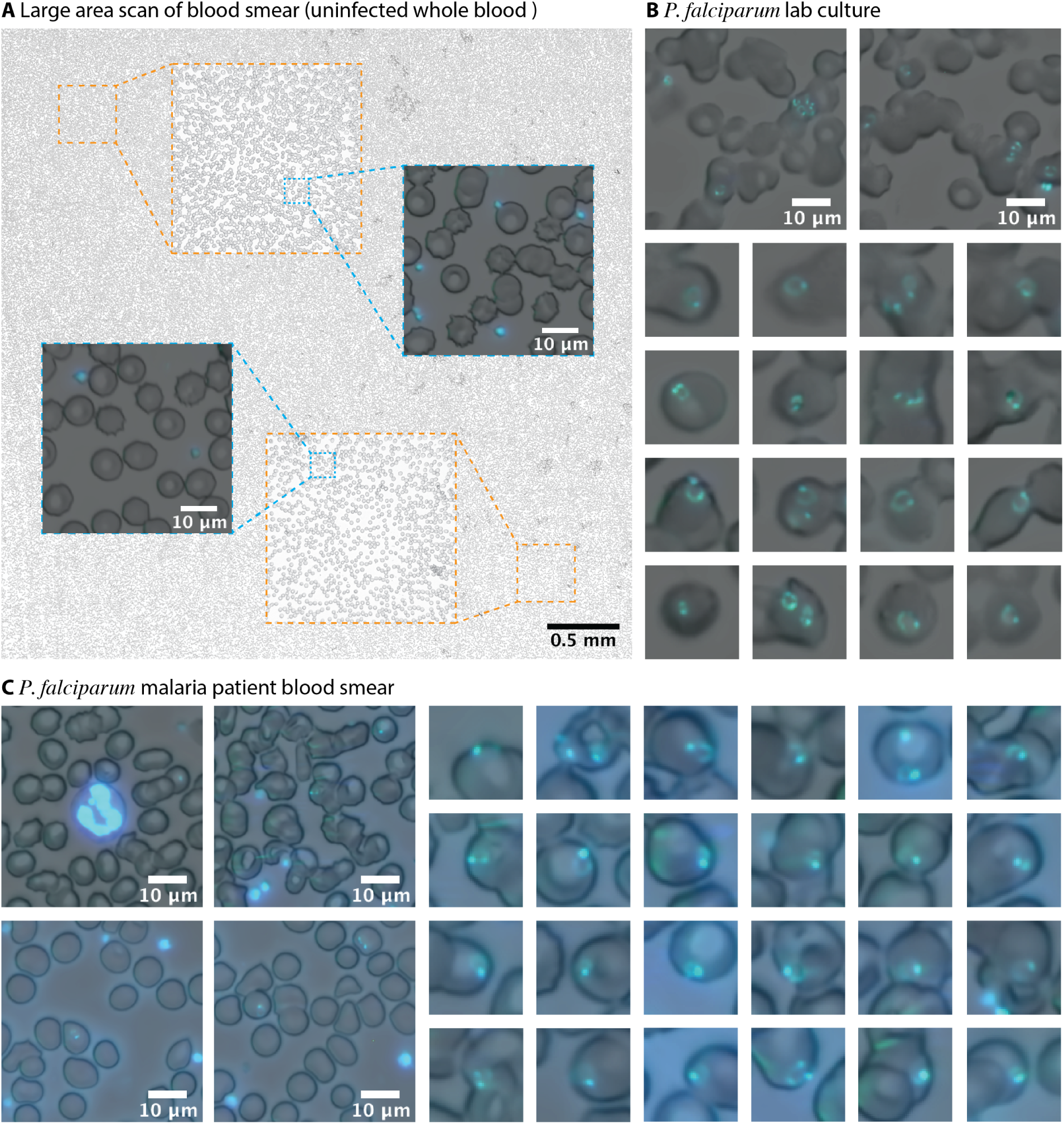
Imaging Blood Smear with the *high mag imaging module*. **(A)** A 11 *×* 11 FOV (4.4mm *×* 4.4mm) scan of DAPI-stained blood smear of a healthy individual. Platelets are visible in the zoomed-in overlaid bright field and fluorescent images **(B)** Selected field of views showing red blood cells infected with ring-stage parasites. Some red blood cells are infected with multiple red blood cells. Ring like morphology of the parasites is clearly visible **(C)** Patient sample showing a white blood cell (top left), platelets and infected red blood cells. Each close-up image in (B) and (C) is of size of 12.9 *µ*m *×* 12.9 *µ*m. Images were denoised by a pretrained FFDNet denosiser [93], see Fig. S13 for images before denoising.

### Broader diagnostics applications

Besides malaria, *Octopi* can be used to image a wide range of pathogens and conditions. As examples, we imaged Schistosomiasis of human intesines tissue specimen (Fig. 8A), *Leishmania donovani* that causes leishmaniasis (Fig. 8B), *Trypanosoma brucei rhodesiense* (Fig. 8C) that causes African sleeping sickness, *Mycobacterium tuberculosis* that causes tuberculosis (TB) (Fig. 8D), *Streptococcus pneumonia* that can cause community-acquired pneumonia (CAP) (Fig. 8E) as well as *Staphylococcus aureus* that can cause bacteremia, skin infection, respiratory infections and food poisoning (Fig. 8F). The last three bacterial pathogens were in sputum samples and imaged using the *high mag imaging module* with a 100x/1.25 oil immersion objective. In the last sample, since the bacteria are distributed in different z-plane, a z-stack was taken to capture all within the field of view.

**Fig. 8.**
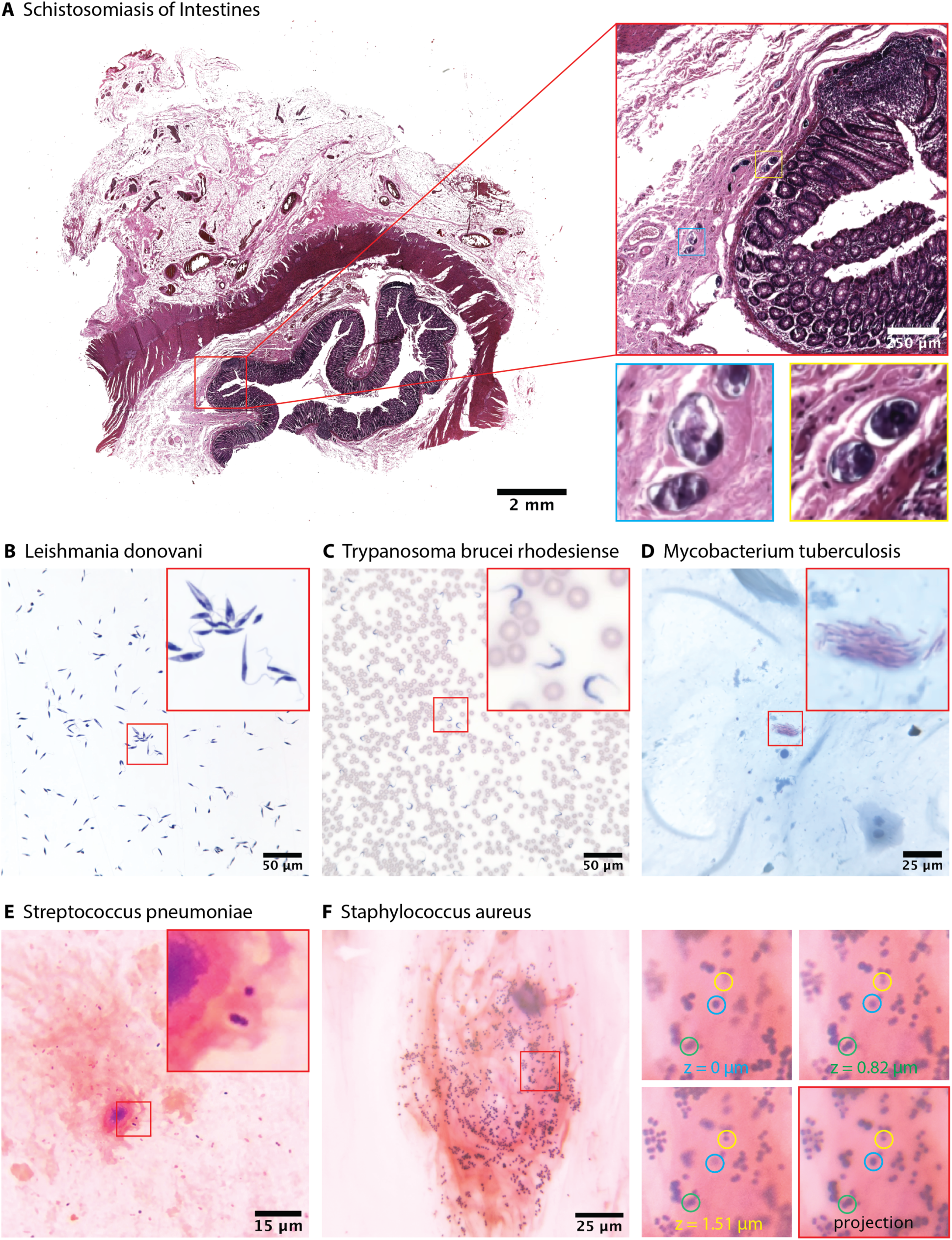
Other diagnostic applications for *Octopi*. **(A)** H&E stained Schistosomiasis of intestines specimen obtained with the *low mag imaging module*. Close-up images show eggs of *Schistosoma haematobium*. **(B)** Hematoxylin stained promastigotes of *Leishmania donovani* obtained with the *high mag imaging module* using a 40x/0.65 Plan Achromatic Objective. **(C)** Giemsa stained *Trypanosoma brucei rhodesiense* in a thin blood smear obtained with the *high mag imaging module* using a 40x/0.65 Plan Achromatic Objective. **(D)** ZiehlNeelsen stained *Mycobacterium tuberculosis* in a sputum sample obtained with the *high mag imaging module* using a 100x/1.25 Plan Achromatic Objective. **(E)** Gram stained *Streptococcus pneumoniae* in a sputum sample obtained with the *high mag imaging module* using a 100x/1.25 Plan Achromatic Objective. **(F)** Gram stained *Staphylococcus aureus* in a sputum sample obtained with the *high mag imaging module* using a 100x/1.25 Plan Achromatic Objective. Left shows minimum intensity projection of a z-stack containing 20 planes with z-step size of 137 nm. Right shows close-up images of different z-planes and the minimum intensity projection corresponding to the same field of view. Images in (A)-(D) were denoised by a pretrained FFDNet denosiser [93], see Fig. S13 for images before denoising.

## Discussion

Here we report the concept and implementation of a modular and automated imaging platform. Compared to directly modifying existing microscopes [56] and many other monolithic designs, the open nature of our platform and its high degree of modularity offers flexibility and greatly simplifies both iterative and derivative developments, making it easy to adapt the tool to specific applications. CNC machining also allows the precision and robustness unmatched by 3D-printing. While we demonstrated bright field, dark field and fluorescent imaging with reversed lens configuration and using standard objectives, other imaging modalities such as Fourier Ptychography [15, 57], holography/lensless imaging [16, 58] as well as standard or LED-matrix and computation-based phase contrast [59, 60] can also be implemented on our platform. Furthermore, metalens made of a single layer of nanostructures [61] may be adopted in place of standard objectives as they become available. The compactness and light weight makes it possible to mount the lens on a voice coil actuator (widely used in cellphone cameras and blu-ray players), eliminating the need for linear stage, which results in reduction in cost and form factor. Metasurfaces or phase masks may also be incorporated into the optical train for aberration correction [62] and for enhanced 3D imaging capability through point spread function engineering [63, 64, 65]. While all images in this report are taken with Pi cameras, CMOS cameras that are low-cost and compact but rival the performance of sCMOS and EMCCD can be used for more demanding applications [66, 67, 68, 69]. With modular design, other XY stage designs may also be used to allow larger travel and/or more precise motion [70, 13, 71].

Applying the imaging platform to the diagnosis of malaria, we developed a new solution that can determine parasitemia with high degree of automation and very high throughput. While there are five strains of parasites that can cause malaria in human, in this study we focused on Plasmodium falciparum (*P. falciparum*) for two reasons. First, *P. falciparum* is the deadliest strain, which can cause fatality if treatment is delayed beyond 24 hours after the onset of clinical symptoms. *P. falciparum* has also developed resistance to nearly all anti-malarials in current use, where chloroquine-resistance has spread to nearly all areas of the world where *P. falciparum* malaria is transmitted [72]. Second, in 2017, the WHO African Region was home to 92% of global malaria cases, out of which 99.7% of is due to *P. falciparum* [73]. The dominance of *P. falciparum* in this region makes our low-mag module-based solution readily applicable without considering the need for speciation. While at low magnification morphology of ring stage parasites cannot be resolved, in the future, other features may be used for speciation. For example, in contrast to *P. falciparum* where most parasites present in the peripheral blood are in ring-stage due to sequestration, both trophozoites and schizonts can be present for P. vivax (Pv). Compared to ring stage, these stages have markedly different morphology features that are likely resolvable even with low magnification [74] and should present more intense and more red-shifted fluorescence.

We have also demonstrated that morphological features of ring-stage parasites can be unambiguously resolved with the *high mag imaging module* on the platform. The *high mag imaging module* may be combined with the *low mag imaging module* for further improved detection limit and speciation capability without sacrificing throughput. To do so, the slide is first screened by the *low mag imaging module*, where locations of suspected pathogens are recorded. The spots are then relocated with the motorized scanner and imaged with the *high mag imaging module* for confirmation. The platform may also be modified to accommodate two imaging modules simultaneously. Encouraged by presented results, we are in planning phase of a clinical trial for testing the efficacy of the instrument in field conditions in both India and Africa.

We have shown that using a color CMOS sensor and long pass emission filter for fluorescence imaging, spectral shift of DAPI-stained *P. falciparum* parasites on the order of 10 nm can be reliably detected in a single shot. This proved to be critical in the application of diagnosis of malaria using blood smears. The loss in spatial resolution compared to using a monochrome sensor under the same condition may be complemented by using higher magnification (without changing the NA of the objective). When the fluorescent spots arise from diffraction limited sources or beads of known size and some emission spectrum information is known a priori, maximum likelihood estimation may be used to optimally extract information. This spectral imaging capability allows single-shot multiplexed detection with a single laser excitation.

While we used nucleic acid stains for sensitive detection of *P. falciparum* parasites in thin blood smear, different probes that are specific to a set of pathogens can also be utilized. The past decades has seen much development of pathogen specific probes [75, 76, 77, 78]. Being low-cost and highly configurable, *Octopi* has the potential to help realize the wide spread use of these new probes in field diagnostics. Besides diagnosis of disease in field conditions, the automated imaging platform can be also be adapted for research applications. In particular, because of its low cost and small footprint, many units can be set up in a single lab to parallelize experiments like super-resolution microscopy [79, 80], expansion microscopy [81], spatially resolved profiling of RNA in single cells [82] and spatial sequencing of single-cell transcriptional states in tissues[83].

With the emergence of deep learning in microscopy, the capabilities of *Octopi* can be boosted by newly developed neural networks that breaks the limits of what is possible in traditional microscopy [84, 85, 86, 87, 88, 84, 89, 90]. As a highly scalable platform, *Octopi* can also help bring training and deployment of these networks to a large number of users. Finally, with a large network of *Octopi* deployed around the world, we envision to bring together researchers, developers and clinical practitioners to collectively advance microscopy-based disease diagnostics.

## Materials and Methods

### Study Design

The goal of this study is to develop and evaluate a low-cost, modular and automated microscope platform for a range of applications including, in particular, diagnosis of infectious disease with high throughput in resource-limited settings. We started by implementing modules of the microscope and characterizing their performances, showing that performance comparable to high end research grade microscope can be achieved. In applying the platform to detection of ring-stage *P. falciparum* parasites, we discovered that with 405 nm laser excitation, a 435 nm long pass emission filter and a color CMOS camera, DAPI-stained parasites and platelets may be told apart by color. We used laser scanning confocal microscopy to obtain spectrum of emitted fluorescence from DAPI stained platelets and parasites in patient sample, which revealed a spectral red-shift on the order of 10 nm. That this shift and the resulting color difference can be used to differentiate parasites and platelets under low magnification was supported by simulation. To automatically detect parasites and quantify diagnostic performance that may be achieved, we collected data from 8 smears of *P. falciparum* culture and 10 smears of uninfected blood and trained a classifier using these data. This amounts to baseline data on 109,3555 spots of parasites (*P. falciparum*) and 437,944 spots for uninfected whole blood. Based on this classifier we simulated sensitivity and specificity that can be expected at different parasitemia. A processing pipeline was implemented on Jetson Nano so that computation can be performed locally in real time. Furthermore, we imaged lab and patient samples on our platform with high magnification to show that morphology of ring-stage parasites can also be resolved, implying further improved sensitivity and specificity can be achieved. Finally, to show broad applicability of our platform, we imaged different prepared pathological samples with different magnifications.

### Construction of the prototype microscope

Custom parts of the microscope were designed with Autodesk Inventor Professional and fabricated by Protolabs and 3D Hubs (CNC machining with 6061 Aluminum), and Fictiv (selective laser sintering with Nylon) (Fig. S9). In the *high mag imaging module*, a piezo stack actuator with end cap (Thorlabs PK2FMP2) was epoxied to the extended contact ball bearing linear stage (SELN LBV40-C2). A 12-bit DAC (Adafruit MCP4725 breakout board) was interfaced with the Raspberry Pi computer through I2C interface. The output of the DAC was amplified by a miniature piezo driver (PiezoDrive PDu100B) to drive the piezo stack actuator. Three stepper motor driver boards (Allegro A3967-based Easydriver) were used to drive the lead screw linear motors and captive linear actuator (Haydon Kerk Pittman 21H4AC-2.5-907).

### Scanning stage flatness characterization

A LabView program was developed to raster scan a target slide (Ossila S151 Ultra-flat Quartz Coated Glass) while recording the relative z-position of top surface of the slide at the center of the microscope field of view, which is measured by a non-contact displacement sensor (MKS Instrument Optimet ConoPoint-3R, Fig. S2). The measurement results were saved as CSV files and processed with MATLAB.

### Deep learning-based red blood cell segmentation

The 91-layer Fully Convolutional DenseNet contains 11 dense blocks (with 4, 5, 7, 10, 12, 15, 12, 10, 7, 5 and 4 layers for each block), and was trained from scratch. Weights of the convolutional layers were initialized using He initialization [91]. For training, Adam optimizer [92] was used with a learning rate of 0.001 and batch size of 16.

To deal with the more frequent false negatives (RBC pixels labeled as non-RBC pixels) compared to false positives (non-RBC pixels as RBC pixels) in the labels of the training data, class weights were introduced in the binary cross-entropy loss function. Specifically, false negatives was associated with a class weight of 10, whereas false positives were associated with a class weight of 0.1. This ensured that mispredictions made on pixels labeled positive, where labels are reliable, are penalized more heavily than mispredictions made on pixels labeled negative, where labels can be noisy.

To obtain a large labeled training data set without tedious human annotation, the following two-step approach is taken. First, Hough transform was used to generate accurate segmentation masks for images where red blood cells are round and isolated. Second, multiple such images were superimposed and distorted through shear transformations to mimic images with red blood cells that are not round and/or overlapping. The resulting images, which also have accurate masks, were used to augment the training data. In total, 22,680 images of size 128 *×* 128 were used for training the neural network.

The benefits of using data augmentation and introduction of class weights are visualized in Fig. S10.

### Spot detection from fluorescent images obtained with the *low mag imaging module*

Two spot detection pipeline were implemented. The CPU-only pipeline (pipeline A) was implemented in python with the scikit-image package. The pipeline that takes advantage of CUDA (pipeline B) was implemented in C++ with the OpenCV library and python with the scikit-image package. Both pipelines take an image of size 1428×1428×3 as input and convert it to grayscale for further processing. In pipeline A, functions skimage.morphology.white tophat and skimage.feature.blob log are used. In pipeline B, image is uploaded to GPU and background removed using tophat filtering with disk diameter of 9. The processed image remains in GPU and is converted from CV U8 to CV F32. Four normalized Laplacian of Gaussian images (LoG1,LoG2,LoG3,LoG4) with gaussian sigam equal to 1, 1.5, 2 and 2.5 are computed by applications of a Gaussian filter followed by a Laplacian filter and scale normalization. The four images are compared with a manually selected threshold, and pixel whose value is smaller than the threshold is set to zero. A maximum projection along the scale dimension of the four LoG images is computed and and a 3×3 maximum filter is applied. The resulting image (P) is compared with the four LoG images, and locations where pixel values equal are recorded in a mask M initialized with zeros (for example, if LoG3(r,c)==P(r,c), then M(r,c) is set to 3). The mask, which stores locations of 3×3×4 local maximums, is downloaded from GPU, and 3D coordinates (2D location + scale) of non-zero elements of the mask are exported as a three column array. The array is loaded in python for removal of spots with overlap exceeding the a set threshold of 0.5. The last step takes advantage of the already implemented skimage.feature.blob. prune blobs function, which uses a KDtree implemented in c to perform nearest neighbour search for significantly reducing the number of pairwise comparison needed.

Before passing to the spot detection processing pipeline, fluorescent images were first converted from sRGB space to linear RGB space, so that pixel intensity has a linear relationship to the number of photons collected. Detected spots were saved for visualization and downstream classification.

### Fluorescent spot classifier

The gradient boosted decision trees classifier was implemented using XGBClassifier from the xgboost python package. Features for each fluorescent spot passed as input to the classifier and their relevance are shown in Fig. S11. Among the features, overlap is the sum of pixel values of pixels that are segmented as part of red blood cells over the sum of pixel values of all pixel. For uninfected whole blood, this feature is directly computed from data. For *P. falciparum* culture, because the red blood cells are ill-shaped for bright field segmentation, this feature was sampled from a empirical distribution, as is plotted in Fig. S12. Performance of the classifier was measured using 20-fold cross validation. Each each fold is made of 3 smears of uninfected whole blood slides and 3 smears of *P. falciparum* lab culture, picked at random. For training the classifier, binary logistic loss function was used with a L2 regularization term.

### Image processing

For all the images presented, image processing were done in MATLAB. For brightfield images, illumination correction is done through normalization against a blank image (image of a blank slide). For fluorescent images, background removal is done through tophat transform. Bright field images from both the *low mag imaging module* and the *high mag imaging module* and fluorescent images from the *high mag imaging module* are demosaiced in MATLAB from the raw bayer data. For images that are denoised (as noted in figure captions), denoising is done using a convolutional neural network FFDNet [93]. Comparisons of images before and after denoising can be found in Fig. S13.

### P. falciparum in vitro cultures

Plasmodium Falciparum culture were provided by the Yeh lab at Stanford University where Plasmodium falciparum W2 (MRA-157) were obtained from MR4. Parasites were grown in human erythrocytes (2% hematocrit, obtained from the Stanford Blood Center) in RPMI 1640 media supplemented with 0.25% Albumax II (GIBCO Life Technologies), 2 g/L sodium bicarbonate, 0.1 mM hypoxanthine, 25 mM HEPES (pH 7.4), and 50 g/L gentamycin, at 37C, 5% O2, and 5% CO2.

### Blood Sample from healthy donors and from patients diagnosed with malaria

De-identified blood sample (whole blood) from healthy anonymous donors were obtained from the Stanford Blood Center in BD Vacutainer blood collection tubes. De-identified methanol-fixed finger prick blood smears from patients diagnosed with malaria were provided by UCSF Malaria Elimination Initiative (MEI)/Infectious Disease Research Collaboration, Kampala, Uganda.

### Preparation and staining of blood smears

Smears of blood from healthy donors and P. falciparum culture were fixed by dipping in absolute methanol for 30 seconds. Fixed smears were incubated with 5 *µ*g/ml DAPI solution for 1 minute, washed in water, and let air dry in the dark. DAPI solution was purchased from Biotium (catalog # 40043) and diluted. Samples were kept in dark before imaging.

### Other pathology samples

Prepared slides of *Loa-Loa*, *Leishmania donovani*, *Mycobacterium tuberculosis* were acquired from VWR (catalog Number 470182-158, 470181-894, 470177-208 respectively). Prepared slide of Schistosomiasis of Intestines was acquired from AmScope (SKU: PS50HP). Prepared slide of Trypanosoma brucei rhodesiense was acquired from Carolina (item # 295822). Prepared slides of *Streptococcus pneumoniae* and *Staphylococcus aureus* in sputum were de-identified and provided by the Clinical Microbiology Laboratory at Stanford Health Care.

### Statistical Analysis

Simulation of per case sensitivity and specificity (Fig. 6E) was done in MATLAB. For each parasitemia and per spot false positive rate (FPR) (and its associated per spot false negative rate FNR), 10,000 tests were simulated. In each test, the total number of platelets (*N*) and parasites (*P*) was sampled from Poisson distributions; the number of detected parasites was the sum of true positives and false positives, both sampled from Bernoulli distribution, with parameters (*P,* 1-FNR) and (*N,* FPR).

## Acknowledgements

We acknowledge Ellen Yeh and her lab including Marta Walczak and Katrina Hong for providing P. falciparum blood cultures. We thank Grant Dorsey and Bryan Greenhouse from UCSF and Harriet Ochokoru, laboratory technician at Infectious Diseases Research Collaboration, Kampala, Uganda for providing de-identified patient blood smears via UCSF Malaria Eliminate program. We thank Brieuc Cossic (Roche), Darvin Scott Smith (Stanford), Niaz Banaei (Stanford) for providing non-malaria samples. We thank Andres Garchitorena and Matt Bonds from PIVOT and Milijaona Randrianarivelojosia from Institute Pasteur, Madagascar and Sister Aquinas and staff of Swasthya Swaraj, Orissa, India and Vasundhara Rangaswamy (Stanford) for fruitful discussions on diagnostics needs in Madagascar and India. We thank MKS Instruments for providing ConoPoint-3 for stage flatness characterization. Finally, we thank Mireille Kamariza (Stanford), Rishabh Manoj Shetty (Arizona State University) and all members of PrakashLab and specifically, Felix Hol, Haripriya Mukundarajan, Scott Coyle, Shailabh Kumar for feedback and comments on the manuscript. **Funding:** H.L. was supported by a Bio-X Stanford Interdisciplinary Graduate Fellowship. L.F.V. was supported by a Stanford Electrical Engineering department fellowship. This research was supported by HHMI-Gates Faculty Fellows Grant (M.P.), NIH New Innovator Award (M.P.), NSF Career Award (M.P.), NSF CCC (grant DBI-1548297) and Moore Foundation. **Author contributions:** H.L. and M.P. designed the instrument and overall research. H.L. built and characterized the instrument. H.L. collected data with assistance from M.P., L.V.F and H.SM.. H.L. developed models for spectral imaging. L.F.V and M.V. developed the neural network for red blood cell segmentation. H.L., L.F.V and M.V. developed processing pipelines and processed the data. H.SM and M.P performed field testing. H.L and M.P wrote the manuscript. **Competing interests:** All authors declare no competing interests. **Data and materials availability:** All data necessary for interpreting the manuscript have been included. Additional information may be requested from the authors.

## List of Supplementary Materials

Fig. S1. CMOS sensor characterization and anticipated performance.

Fig. S2. Setup for scanning stage flatness characterization.

Fig. S3. Characterization of XY scanning flatness for scanning module using a normal microscope glass slide.

Fig. S4. Per frame processing time of Top-hat transform and LoG Spot detection implemented on different computation modules.

Fig. S5. Illustration of the spot detection pipeline.

Fig. S6. Deep learning red blood cell segmentation.

Fig. S7. Full field of view pseudo color image of uninfected whole blood.

Fig. S8. Full field of view pseudo color image of blood smear from patient diagnosed of *P. falciparum* malaria.

Fig. S9. Parts common to different configurations of *octopi* presented.

Fig. S10. Training of the convolutional neural network.

Fig. S11. Features used by the classifier and their importance scores output by the classifier in the training phase.

Fig. S12. Empirical distribution of overlap between red blood cells and *P. falciparum* parasites

Fig. S13. Images before and after denoising.

Table S1. Cost breakdown for *Octopi*.

Movie S1. Operation of the *Octopi* platform for automated slide scanning.

Movie S2. Focus adjustment with Piezo actuator while imaging a resolution target.

## References

1. E. Yusuf and R. L. Hamers, “What the WHO’s List of Essential Diagnostics means for clinical microbiology laboratories and antimicrobial stewardship practice worldwide,” Clinical Microbiology and Infection, vol. 25, pp. 6–9, jan 2019.

2. E. Fu, P. Yager, P. N. Floriano, N. Christodoulides, and J. T. McDevitt, “Perspective on diagnostics for global health,” IEEE Pulse, vol. 2, no. 6, pp. 40–50, 2011.

3. K. J. Land, D. I. Boeras, X.-S. Chen, A. R. Ramsay, and R. W. Peeling, “Reassured diagnostics to inform disease control strategies, strengthen health systems and improve patient outcomes,” Nature microbiology, vol. 4, no. 1, p. 46, 2019.

4. D. Berger, “A brief history of medical diagnosis and the birth of the clinical laboratory. Part 1–Ancient times through the 19th century.,” MLO: medical laboratory observer, vol. 31, pp. 28–30, 32, 34–40, jul 1999.

5. B. A. Mathison and B. S. Pritt, “Update on Malaria Diagnostics and Test Utilization,” Journal of Clinical Microbiology, vol. 55, no. 7, pp. 2009–2017, 2017.

6. WHO, “Malaria microscopy quality assurance manual Ver. 2,” tech. rep., 2016.

7. N. Farahani and C. E. Monteith, “The coming paradigm shift: A transition from manual to automated microscopy,” Journal of pathology informatics, vol. 7, p. 35, sep 2016.

8. A. F. Coskun, S. N. Topkaya, A. K. Yetisen, and A. E. Cetin, “Portable multiplex optical assays,” Advanced Optical Materials, vol. 7, no. 4, p. 1801109, 2019.

9. Q. Wei, H. Qi, W. Luo, D. Tseng, S. J. Ki, Z. Wan, Z. Grcs, L. A. Bentolila, T.-T. Wu, R. Sun, et al., “Fluorescent imaging of single nanoparticles and viruses on a smart phone,” ACS nano, vol. 7, no. 10, pp. 9147–9155, 2013.

10. J. S. Cybulski, J. Clements, and M. Prakash, “Foldscope: origami-based paper microscope,” PloS one, vol. 9, no. 6, p. e98781, 2014.

11. A. Skandarajah, C. D. Reber, N. A. Switz, and D. A. Fletcher, “Quantitative imaging with a mobile phone microscope,” PloS one, vol. 9, no. 5, p. e96906, 2014.

12. M. V. DAmbrosio, M. Bakalar, S. Bennuru, C. Reber, A. Skandarajah, L. Nilsson, N. Switz, J. Kamgno, S. Pion, M. Boussinesq, et al., “Point-of-care quantification of blood-borne filarial parasites with a mobile phone microscope,” Science translational medicine, vol. 7, no. 286, pp. 286re4–286re4, 2015.

13. J. P. Sharkey, D. C. Foo, A. Kabla, J. J. Baumberg, and R. W. Bowman, “A one-piece 3d printed flexure translation stage for open-source microscopy,” Review of Scientific Instruments, vol. 87, no. 2, p. 025104, 2016.

14. O. Holmström, N. Linder, B. Ngasala, A. Mårtensson, E. Linder, M. Lundin, H. Moilanen, A. Suutala, V. Diwan, and J. Lundin, “Point-of-care mobile digital microscopy and deep learning for the detection of soil-transmitted helminths and schistosoma haematobium,” Global health action, vol. 10, no. sup3, p. 1337325, 2017.

15. G. Zheng, R. Horstmeyer, and C. Yang, “Wide-field, high-resolution Fourier ptychographic microscopy,” Nature Photonics, vol. 7, no. 9, pp. 739–745, 2013.

16. A. Greenbaum, Y. Zhang, A. Feizi, P.-L. Chung, W. Luo, S. R. Kandukuri, and A. Ozcan, “Wide-field computational imaging of pathology slides using lens-free on-chip microscopy,” Science translational medicine, vol. 6, no. 267, pp. 267ra175–267ra175, 2014.

17. A. Ozcan and E. McLeod, “Lensless imaging and sensing,” Annual review of biomedical engineering, vol. 18, pp. 77–102, 2016.

18. Y. Wu and A. Ozcan, “Lensless digital holographic microscopy and its applications in biomedicine and environmental monitoring,” Methods, vol. 136, pp. 4–16, 2018.

19. A. Ozcan, “Mobile phones democratize and cultivate next-generation imaging, diagnostics and measurement tools,” Lab on a Chip, vol. 14, no. 17, pp. 3187–3194, 2014.

20. I. Hernández-Neuta, F. Neumann, J. Brightmeyer, T. Ba Tis, N. Madaboosi, Q. Wei, A. Ozcan, and M. Nilsson, “Smartphone-based clinical diagnostics: towards democratization of evidence-based health care,” Journal of internal medicine, vol. 285, no. 1, pp. 19–39, 2019.

21. WHO, “World Malaria Report. 2018. ISBN 978 92 4 156469 4.,” 2018.

22. J. Daily, “Malaria diagnostics technology and market landscape,” 2016.

23. U. Dalrymple, R. Arambepola, P. W. Gething, and E. Cameron, “How long do rapid diagnostic tests remain positive after anti-malarial treatment?,” Malaria journal, vol. 17, no. 1, p. 228, 2018.

24. D. Gamboa, M.-F. Ho, J. Bendezu, K. Torres, P. L. Chiodini, J. W. Barnwell, S. Incardona, M. Perkins, D. Bell, J. McCarthy, et al., “A large proportion of p. falciparum isolates in the amazon region of peru lack pfhrp2 and pfhrp3: implications for malaria rapid diagnostic tests,” PloS one, vol. 5, no. 1, p. e8091, 2010.

25. W. H. Organization, et al., “False-negative rdt results and implications of new reports of p. falciparum histidine-rich protein 2/3 gene deletions,” tech. rep., World Health Organization, 2017.

26. C. L. Cohen, Y. Eshel, J. Swaminathan, D. Gluck, D. Maina, C. Mbithi, S. Onsongo, Z. Premji, N. Lezmy, Z. Nneka, S. Levy-Schreier, J. J. Pollak, S. J. Salpeter, H. Benkuzari, H. Solomon, M. Charles, P. Sampathkumar, M. Soni, and A. Houri-Yafin, “Evaluation of the Parasight Platform for Malaria Diagnosis,” Journal of Clinical Microbiology, vol. 55, no. 3, pp. 768–775, 2017.

27. C. B. Delahunt, C. Mehanian, L. Hu, S. K. McGuire, C. R. Champlin, M. P. Horning, B. K. Wilson, and C. M. Thompon, “Automated microscopy and machine learning for expert-level malaria field diagnosis,” in 2015 IEEE Global Humanitarian Technology Conference (GHTC), pp. 393–399, IEEE, 2015.

28. A. Maxmen, “How to defuse malaria’s ticking time bomb,” Nature, vol. 559, pp. 458– 465, 2018.

29. K. Haldar, S. Bhattacharjee, and I. Safeukui, “Drug resistance in Plasmodium,” Nature Reviews Microbiology, vol. 16, no. 3, pp. 156–170, 2018.

30. A. J. Abdifatah, C. Wanna, and N.-B. Kesara, “Plasmodium falciparum drug resistance gene status in the Horn of Africa: A systematic review,” African Journal of Pharmacy and Pharmacology, vol. 12, no. 25, pp. 361–373, 2018.

31. WHO, “Status report on artemisinin resistance and ACT efficacy (August 2018),” no. August, 2018.

32. W. H. Organization, Malaria Microscopy Quality Assurance Manual-Version 2. World Health Organization, 2016.

33. N. A. Switz, M. V. D’Ambrosio, and D. A. Fletcher, “Low-cost mobile phone microscopy with a reversed mobile phone camera lens,” PloS one, vol. 9, no. 5, p. e95330, 2014.

34. S. Jiang, Z. Bian, X. Huang, P. Song, H. Zhang, Y. Zhang, and G. Zheng, “Rapid and robust whole slide imaging based on led-array illumination and color-multiplexed single-shot autofocusing,” arXiv preprint arXiv:1905.03371, 2019.

35. H. Pinkard, Z. Phillips, A. Babakhani, D. A. Fletcher, and L. Waller, “Deep learning for single-shot autofocus microscopy,” Optica, vol. 6, no. 6, pp. 794–797, 2019.

36. J. Sun, C. Zuo, J. Zhang, Y. Fan, and Q. Chen, “High-speed Fourier ptychographic microscopy based on programmable annular illuminations,” Scientific Reports, vol. 8, no. 1, pp. 1–12, 2018.

37. A. GmbH, “Dapi, product no. a1001,”

38. F. Otto, “Dapi staining of fixed cells for high-resolution flow cytometry of nuclear dna,” in Methods in cell biology, vol. 33, pp. 105–110, Elsevier, 1990.

39. T. Lindeberg, “Scale-space theory: A basic tool for analyzing structures at different scales,” Journal of applied statistics, vol. 21, no. 1-2, pp. 225–270, 1994.

40. T. Lindeberg, “Feature detection with automatic scale selection,” International journal of computer vision, vol. 30, no. 2, pp. 79–116, 1998.

41. S. Jégou, M. Drozdzal, D. Vazquez, A. Romero, and Y. Bengio, “The one hundred layers tiramisu: Fully convolutional densenets for semantic segmentation,” in Proceedings of the IEEE Conference on Computer Vision and Pattern Recognition Workshops, pp. 11– 19, 2017.

42. S. Han, H. Mao, and W. J. Dally, “Deep compression: Compressing deep neural networks with pruning, trained quantization and huffman coding,” arXiv preprint arXiv:1510.00149, 2015.

43. L.-C. Chen, Y. Zhu, G. Papandreou, F. Schroff, and H. Adam, “Encoder-decoder with atrous separable convolution for semantic image segmentation,” in ECCV, 2018.

44. M. Sandler, A. Howard, M. Zhu, A. Zhmoginov, and L.-C. Chen, “Mobilenetv2: Inverted residuals and linear bottlenecks,” in CVPR, 2018.

45. B. Sun, L. Yang, P. Dong, W. Zhang, J. Dong, and C. Young, “Ultra power-efficient cnn domain specific accelerator with 9.3tops/watt for mobile and embedded applications,” in The IEEE Conference on Computer Vision and Pattern Recognition (CVPR) Workshops, June 2018.

46. F. Kawamoto, “Rapid diagnosis of malaria by fluorescence microscopy with light microscope and interference filter,” The Lancet, vol. 337, no. 8735, pp. 200–202, 1991.

47. R. Guy, P. Liu, P. Pennefather, and I. Crandall, “The use of fluorescence enhancement to improve the microscopic diagnosis of falciparum malaria,” Malaria journal, vol. 6, no. 1, p. 89, 2007.

48. M. P. Horning, C. B. Delahunt, S. R. Singh, S. H. Garing, and K. P. Nichols, “A paper microfluidic cartridge for automated staining of malaria parasites with an optically transparent microscopy window,” Lab on a chip, vol. 14, no. 12, pp. 2040–2046, 2014.

49. J. Kapuscinski, “Interactions of nucleic acids with fluorescent dyes: spectral properties of condensed complexes.,” Journal of Histochemistry & Cytochemistry, vol. 38, no. 9, pp. 1323–1329, 1990.

50. S. H. Apte, P. L. Groves, J. S. Roddick, V. P. da Hora, and D. L. Doolan, “High-throughput multi-parameter flow-cytometric analysis from micro-quantities of plasmodium-infected blood,” International journal for parasitology, vol. 41, no. 12, pp. 1285–1294, 2011.

51. S. Cervantes, J. Prudhomme, D. Carter, K. G. Gopi, Q. Li, Y.-T. Chang, and K. G. Le Roch, “High-content live cell imaging with rna probes: advancements in high-throughput antimalarial drug discovery,” BMC cell biology, vol. 10, no. 1, p. 45, 2009.

52. N. Regev-Rudzki, D. W. Wilson, T. G. Carvalho, X. Sisquella, B. M. Coleman, M. Rug, D. Bursac, F. Angrisano, M. Gee, A. F. Hill, et al., “Cell-cell communication between malaria-infected red blood cells via exosome-like vesicles,” Cell, vol. 153, no. 5, pp. 1120–1133, 2013.

53. X. Sisquella, Y. Ofir-Birin, M. A. Pimentel, L. Cheng, P. A. Karam, N. G. Sampaio, J. S. Penington, D. Connolly, T. Giladi, B. J. Scicluna, et al., “Malaria parasite dna-harbouring vesicles activate cytosolic immune sensors,” Nature communications, vol. 8, no. 1, p. 1985, 2017.

54. K. A. Babatunde, S. Mbagwu, M. A. Hernández-Castañeda, S. R. Adapa, M. Walch, L. Filgueira, L. Falquet, R. H. Jiang, I. Ghiran, and P.-Y. Mantel, “Malaria infected red blood cells release small regulatory rnas through extracellular vesicles,” Scientific reports, vol. 8, no. 1, p. 884, 2018.

55. Y. Ofir-Birin, P. Abou Karam, A. Rudik, T. Giladi, Z. Porat, and N. Regev-Rudzki, “Monitoring extracellular vesicle cargo active uptake by imaging flow cytometry,” Frontiers in immunology, vol. 9, p. 1011, 2018.

56. Z. Bian, G. Zheng, K. Guo, X. Heng, and J. Liao, “InstantScope: a low-cost whole slide imaging system with instant focal plane detection,” Biomedical Optics Express, vol. 6, no. 9, p. 3210, 2015.

57. T. Aidukas, R. Eckert, A. R. Harvey, L. Waller, and P. C. Konda, “Low-cost, sub-micron resolution, wide-field computational microscopy using opensource hardware,” Scientific reports, vol. 9, no. 1, p. 7457, 2019.

58. N. Antipa, G. Kuo, R. Heckel, B. Mildenhall, E. Bostan, R. Ng, and L. Waller, “Diffusercam: lensless single-exposure 3d imaging,” Optica, vol. 5, no. 1, pp. 1–9, 2018.

59. L. Tian, J. Wang, and L. Waller, “3d differential phase-contrast microscopy with computational illumination using an led array,” Optics letters, vol. 39, no. 5, pp. 1326–1329, 2014.

60. L. Tian and L. Waller, “Quantitative differential phase contrast imaging in an led array microscope,” Optics express, vol. 23, no. 9, pp. 11394–11403, 2015.

61. M. Khorasaninejad, W. T. Chen, R. C. Devlin, J. Oh, A. Y. Zhu, and F. Capasso, “Metalenses at visible wavelengths: Diffraction-limited focusing and subwavelength resolution imaging,” Science, vol. 352, no. 6290, pp. 1190–1194, 2016.

62. W. T. Chen, A. Y. Zhu, J. Sisler, Y.-W. Huang, K. M. Yousef, E. Lee, C.-W. Qiu, and F. Capasso, “Broadband achromatic metasurface-refractive optics,” Nano letters, vol. 18, no. 12, pp. 7801–7808, 2018.

63. S. R. P. Pavani, M. A. Thompson, J. S. Biteen, S. J. Lord, N. Liu, R. J. Twieg, R. Piestun, and W. Moerner, “Three-dimensional, single-molecule fluorescence imaging beyond the diffraction limit by using a double-helix point spread function,” Proceedings of the National Academy of Sciences, vol. 106, no. 9, pp. 2995–2999, 2009.

64. Y. Shechtman, S. J. Sahl, A. S. Backer, and W. Moerner, “Optimal point spread function design for 3d imaging,” Physical review letters, vol. 113, no. 13, p. 133902, 2014.

65. A. S. Backer and W. Moerner, “Extending single-molecule microscopy using optical fourier processing,” The Journal of Physical Chemistry B, vol. 118, no. 28, pp. 8313– 8329, 2014.

66. H. Ma, R. Fu, J. Xu, and Y. Liu, “A simple and cost-effective setup for super-resolution localization microscopy,” Scientific reports, vol. 7, no. 1, p. 1542, 2017.

67. R. Diekmann, K. Till, M. Müller, M. Simonis, M. Schüttpelz, and T. Huser, “Characterization of an industry-grade cmos camera well suited for single molecule localization microscopy–high performance super-resolution at low cost,” Scientific reports, vol. 7, no. 1, p. 14425, 2017.

68. H. P. Babcock, “Multiplane and spectrally-resolved single molecule localization microscopy with industrial grade cmos cameras,” Scientific reports, vol. 8, no. 1, p. 1726, 2018.

69. B. Diederich, P. Then, A. Jügler, R. Förster, and R. Heintzmann, “cellstormcost-effective super-resolution on a cellphone using dstorm,” PloS one, vol. 14, no. 1, p. e0209827, 2019.

70. R. A. Campbell, R. W. Eifert, and G. C. Turner, “Openstage: a low-cost motorized microscope stage with sub-micron positioning accuracy,” PloS one, vol. 9, no. 2, p. e88977, 2014.

71. D. Schneidereit, L. Kraus, J. C. Meier, O. Friedrich, and D. F. Gilbert, “Step-by-step guide to building an inexpensive 3d printed motorized positioning stage for automated high-content screening microscopy,” Biosensors and Bioelectronics, vol. 92, pp. 472–481, 2017.

72. P. B. Bloland, W. H. Organization, et al., “Drug resistance in malaria,” tech. rep., Geneva: World Health Organization, 2001.

73. W. H. Organization, World malaria report 2018. World Health Organization, 2019.

74. W. H. Organization and C. for Disease Control, Basic malaria microscopy. World Health Organization, 2010.

75. E. F. DeLong, G. S. Wickham, and N. R. Pace, “Phylogenetic stains: ribosomal rna-based probes for the identification of single cells,” Science, vol. 243, no. 4896, pp. 1360– 1363, 1989.

76. J. Shah, O. Mark, H. Weltman, N. Barcelo, W. Lo, D. Wronska, S. Kakkilaya, A. Rao, S. T. Bhat, R. Sinha, et al., “Fluorescence in situ hybridization (fish) assays for diagnosing malaria in endemic areas,” PLoS One, vol. 10, no. 9, p. e0136726, 2015.

77. M. Kamariza, P. Shieh, C. S. Ealand, J. S. Peters, B. Chu, F. P. Rodriguez-Rivera, M. R. B. Sait, W. V. Treuren, N. Martinson, R. Kalscheuer, et al., “Rapid detection of mycobacterium tuberculosis in sputum with a solvatochromic trehalose probe,” Science translational medicine, vol. 10, no. 430, p. eaam6310, 2018.

78. Y. Cheng, J. Xie, K.-H. Lee, R. L. Gaur, A. Song, T. Dai, H. Ren, J. Wu, Z. Sun, N. Banaei, et al., “Rapid and specific labeling of single live mycobacterium tuberculosis with a dual-targeting fluorogenic probe,” Science translational medicine, vol. 10, no. 454, p. eaar4470, 2018.

79. E. Betzig, G. H. Patterson, R. Sougrat, O. W. Lindwasser, S. Olenych, J. S. Bonifacino, M. W. Davidson, J. Lippincott-Schwartz, and H. F. Hess, “Imaging intracellular fluorescent proteins at nanometer resolution,” Science, vol. 313, no. 5793, pp. 1642–1645, 2006.

80. M. J. Rust, M. Bates, and X. Zhuang, “Sub-diffraction-limit imaging by stochastic optical reconstruction microscopy (storm),” Nature methods, vol. 3, no. 10, p. 793, 2006.

81. F. Chen, P. W. Tillberg, and E. S. Boyden, “Expansion microscopy,” Science, vol. 347, no. 6221, pp. 543–548, 2015.

82. K. H. Chen, A. N. Boettiger, J. R. Moffitt, S. Wang, and X. Zhuang, “Spatially resolved, highly multiplexed rna profiling in single cells,” Science, vol. 348, no. 6233, p. aaa6090, 2015.

83. X. Wang, W. E. Allen, M. A. Wright, E. L. Sylwestrak, N. Samusik, S. Vesuna, K. Evans, C. Liu, C. Ramakrishnan, J. Liu, et al., “Three-dimensional intact-tissue sequencing of single-cell transcriptional states,” Science, vol. 361, no. 6400, p. eaat5691, 2018.

84. M. Weigert, U. Schmidt, T. Boothe, A. Müller, A. Dibrov, A. Jain, B. Wilhelm, D. Schmidt, C. Broaddus, S. Culley, et al., “Content-aware image restoration: pushing the limits of fluorescence microscopy,” Nature methods, vol. 15, no. 12, p. 1090, 2018.

85. Y. Rivenson, Z. Göröcs, H. Günaydin, Y. Zhang, H. Wang, and A. Ozcan, “Deep learning microscopy,” Optica, vol. 4, no. 11, pp. 1437–1443, 2017.

86. Y. Rivenson and A. Ozcan, “Toward a thinking microscope: Deep learning in optical microscopy and image reconstruction,” arXiv preprint arXiv:1805.08970, 2018.

87. Y. Rivenson, H. Ceylan Koydemir, H. Wang, Z. Wei, Z. Ren, H. Gnaydın, Y. Zhang, Z. Gorocs, K. Liang, D. Tseng, et al., “Deep learning enhanced mobile-phone microscopy,” ACS Photonics, vol. 5, no. 6, pp. 2354–2364, 2018.

88. H. Wang, Y. Rivenson, Y. Jin, Z. Wei, R. Gao, H. Günaydın, L. A. Bentolila, C. Kural, and A. Ozcan, “Deep learning enables cross-modality super-resolution in fluorescence microscopy,” Nat. Methods, vol. 16, pp. 103–110, 2019.

89. Y. Rivenson, T. Liu, Z. Wei, Y. Zhang, K. de Haan, and A. Ozcan, “Phasestain: the digital staining of label-free quantitative phase microscopy images using deep learning,” Light: Science & Applications, vol. 8, no. 1, p. 23, 2019.

90. Y. Rivenson, H. Wang, Z. Wei, K. de Haan, Y. Zhang, Y. Wu, H. Günaydın, J. E. Zuckerman, T. Chong, A. E. Sisk, et al., “Virtual histological staining of unlabelled tissue-autofluorescence images via deep learning,” Nature Biomedical Engineering, p. 1, 2019.

91. K. He, X. Zhang, S. Ren, and J. Sun, “Delving deep into rectifiers: Surpassing human-level performance on imagenet classification,” in Proceedings of the IEEE international conference on computer vision, pp. 1026–1034, 2015.

92. D. P. Kingma and J. Ba, “Adam: A method for stochastic optimization,” arXiv preprint arXiv:1412.6980, 2014.

93. K. Zhang, W. Zuo, and L. Zhang, “Ffdnet: Toward a fast and flexible solution for cnn-based image denoising,” IEEE Transactions on Image Processing, vol. 27, no. 9, pp. 4608–4622, 2018.

