## Supplementary Figures and Tables for "Octopi: Open configurable high-throughput imaging platform for infectious disease diagnosis in the field"

June 26, 2019

### Supplementary Figures

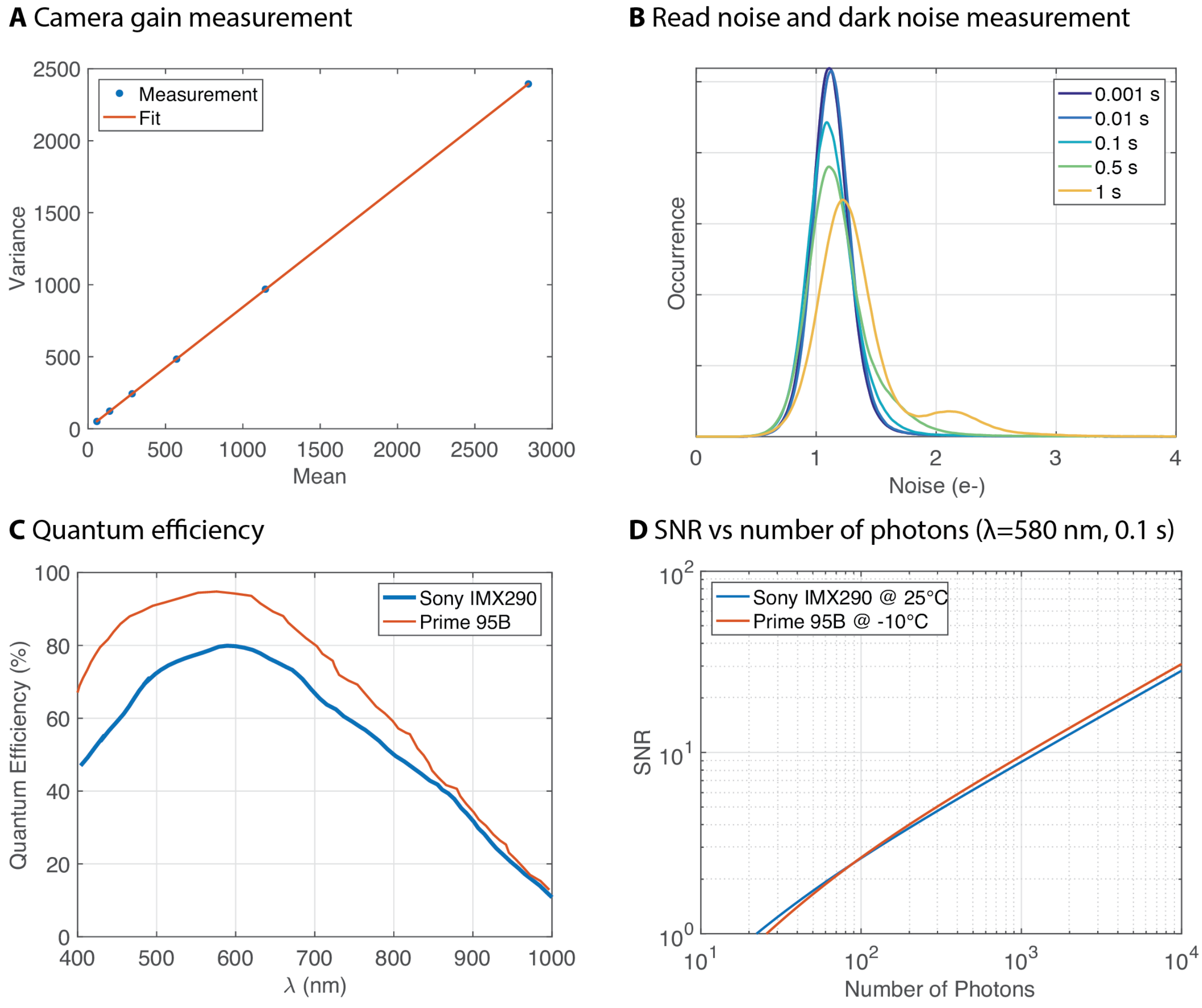

**Figure S1** CMOS sensor characterization and anticipated performance. **(A)** Mean-Variance plot obtained at camera analog gain of 10 dB. Fit of the curve is used to determine the photo-electron analog digital unit (ADU) conversion factor, using which conversion gain at different camera analog gain can be calculated. **(B)** Distribution of pixel value standard deviation of dark image stacks of different exposure time (100 images each, analog gain = 20 dB). The peak reflects the noise level of combined readout noise and noise originating from dark current from the majority of the pixels. The second peak for the 1 s exposure time is from the population of hot pixels. **(C)** Quantum efficiency for different sensors. Data are obtained from sensor datasheets. **(D)** Calculated SNR (to be expected) for different sensors at different signal level (number of photons), assuming 1 s exposure time and wavelength of 850 nm.

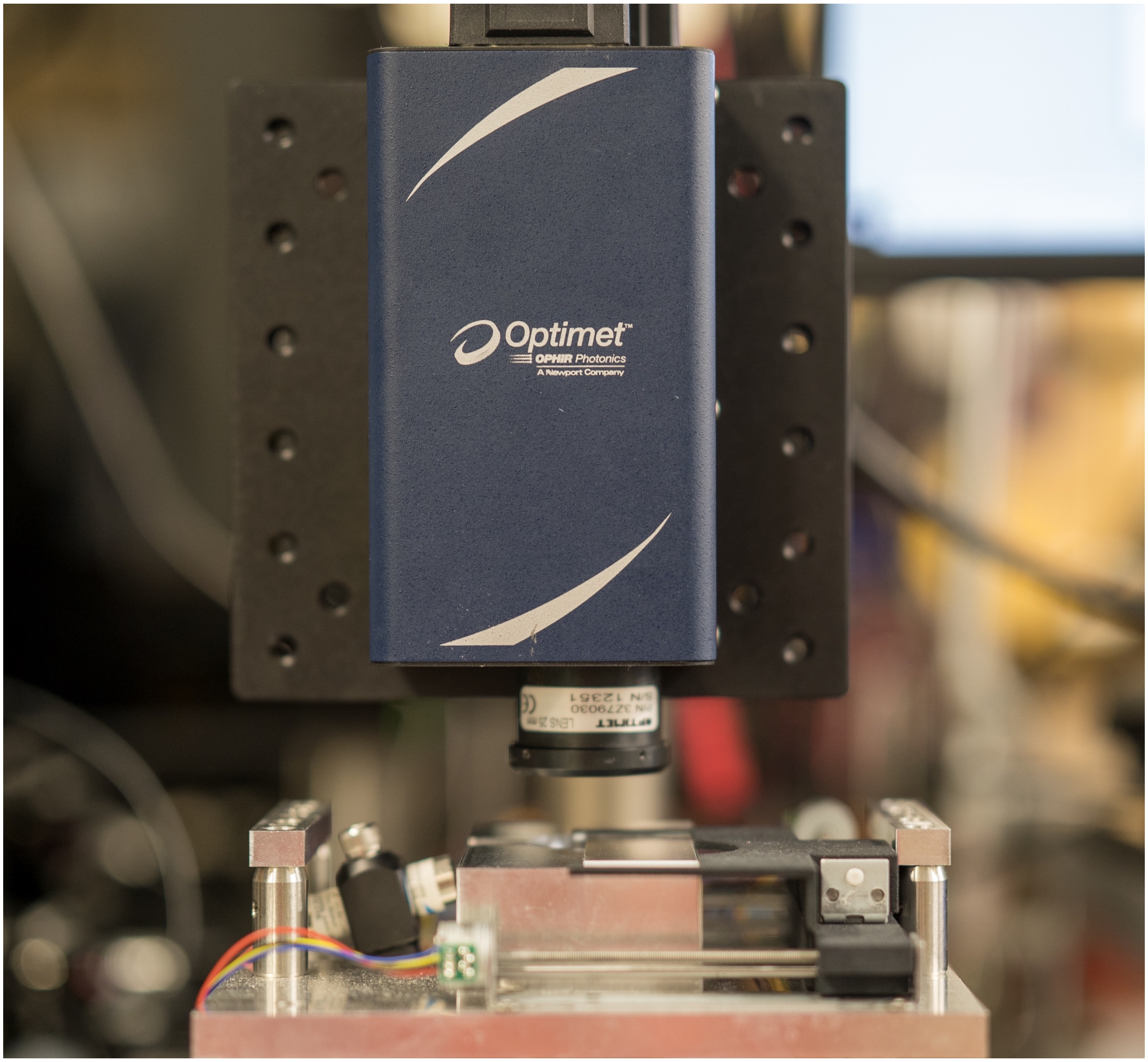

**Figure S2** Setup for scanning stage flatness characterization. The relative z position of the target slide at the center of the platform is measured with a optical conoscopic holography sensor (MKS Instrument Optimet ConoPoint-3R).

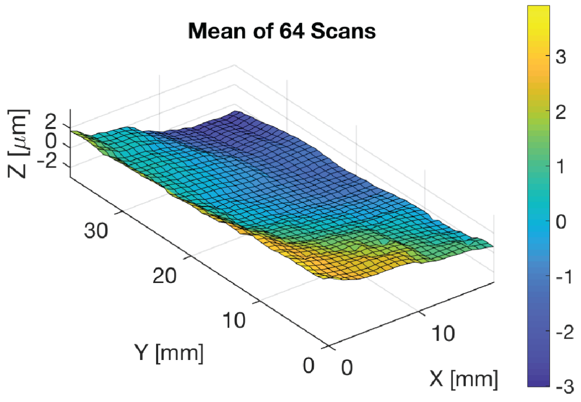

**Figure S3** Characterization of XY scanning flatness for scanning module using a normal microscope glass slide. Mean and standard deviation of measured top surface z-positions for 64 XY scans are plotted. The overall standard deviation is below 400 nm and is limited by the measurement setup, suggesting excellent stage flatness.

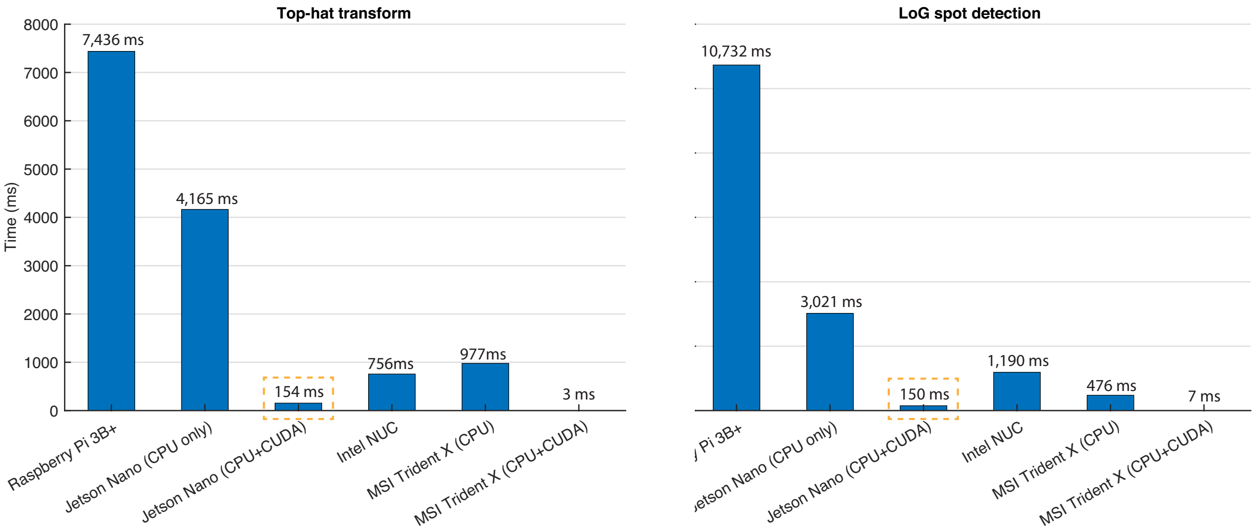

**Figure S4** Per frame processing time of Top-hat transform and LoG Spot detection implemented on different computation modules. Intel NUC is model NUC8i7BEH with Intel Core i7-8559U processor. MSI Trident X is model 9SD-055US with Intel Core i7-9700K processor and GeForce RTX 2070 ARMOR 8G graphics card. We note that Jetson Nano with CUDA implementation is more than 50 times faster than Raspberry Pi 3B+

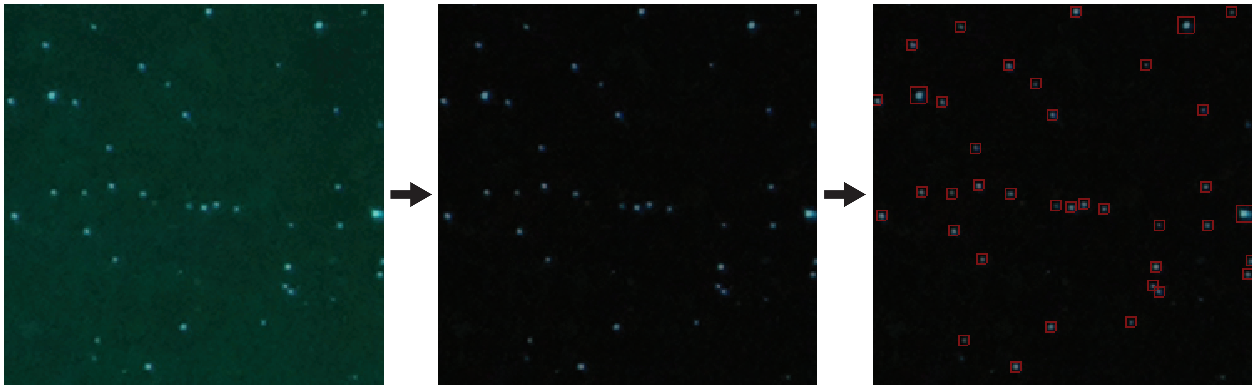

**Figure S5** Illustration of the spot detection pipeline. The image is first background removed with top-hat transform. Spots are detected (shown inside red bounding boxes) using a Laplacian of Gaussian-based detector.

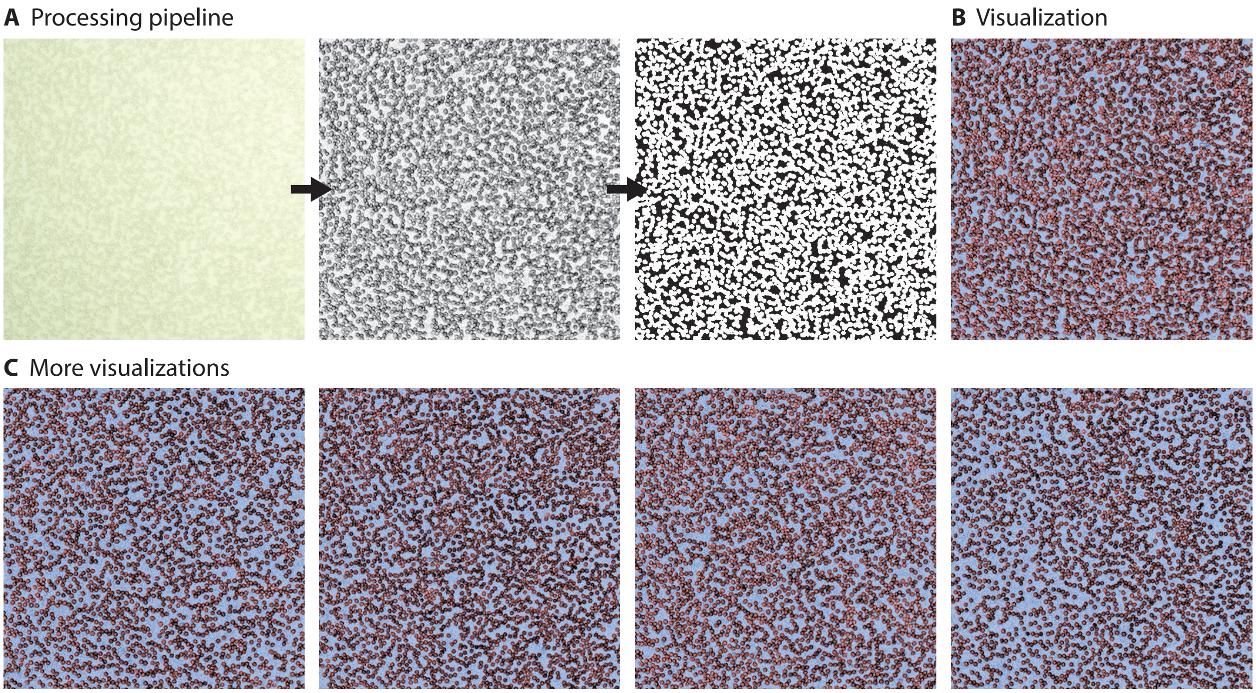

**Figure S6** Deep learning red blood cell segmentation. **(A)** Processing pipeline. The bright-field image is first converted to grayscale, illumination corrected and contrast adjusted. The resulting image is fed to a neural network, which outputs a segmentation mask. **(B)** Processed bright-field image with segmentation mask overlaid. **(C)** Processed bright-field images with segmentation masks overlaid for four more fields of view.

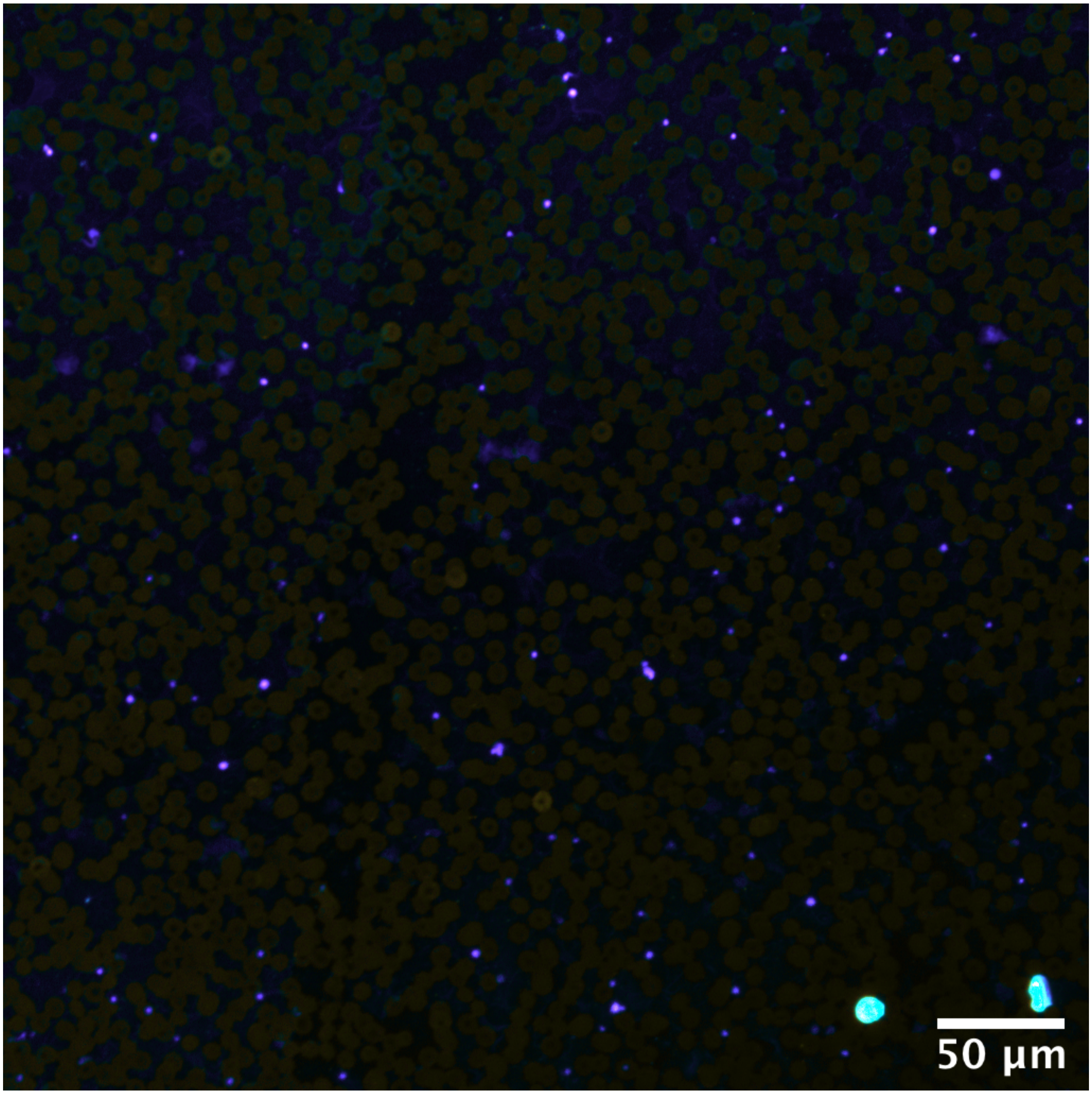

**Figure S7** Full field of view pseudo color image of uninfected whole blood. The image is obtained with laser scanning confocal microscopy with 32 spectral channels. Color of each pixel is determined by the centroid of the spectrum at that pixel. Colorbar is plotted in Fig. 4B

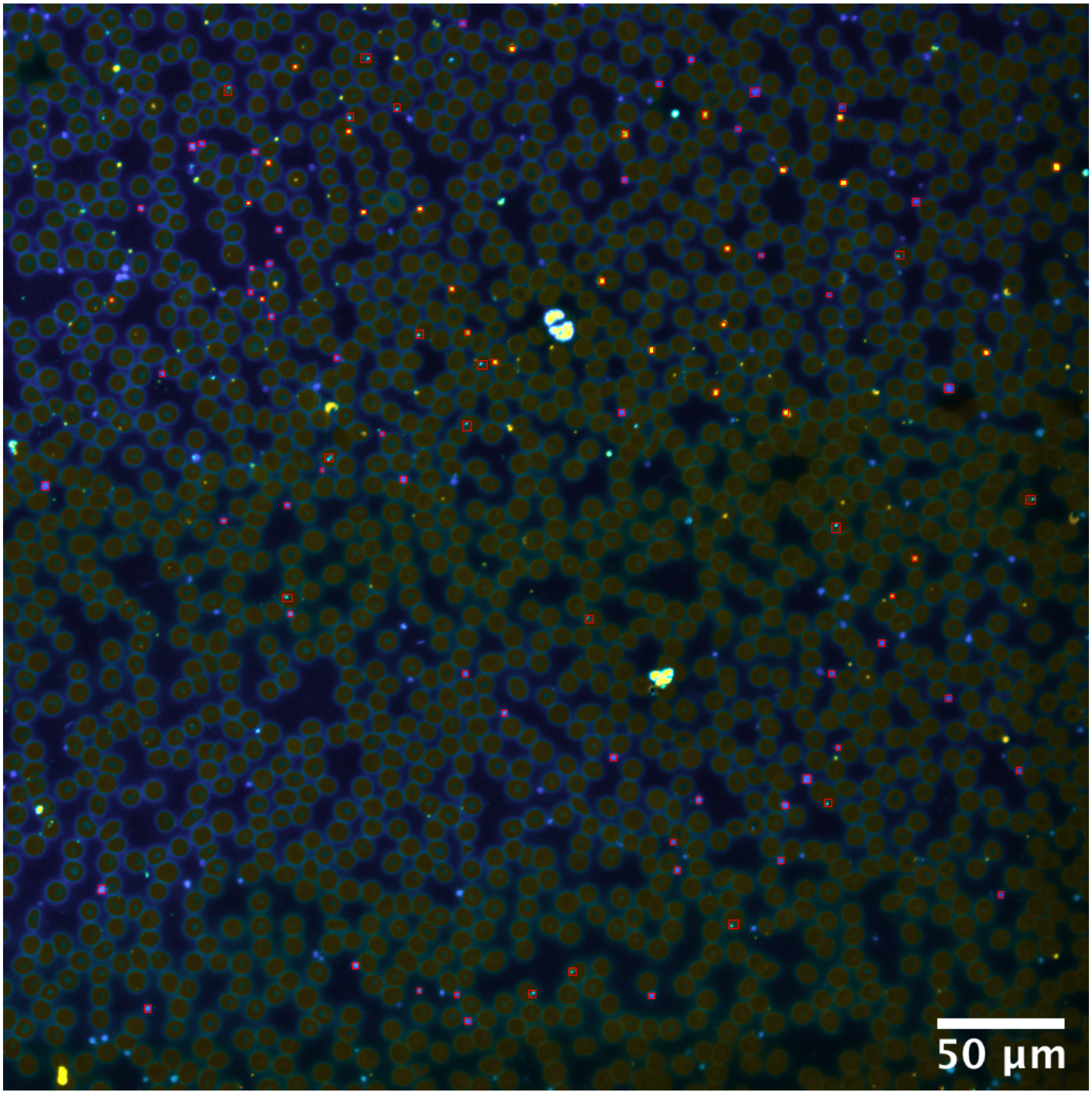

**Figure S8** Full field of view pseudo color image of blood smear from patient diagnosed of *P. falciparum* malaria. The image is obtained with laser scanning confocal microscopy with 32 spectral channels. Color of each pixel is determined by the centroid of the spectrum at that pixel. Colorbar is plotted in Fig. 4B

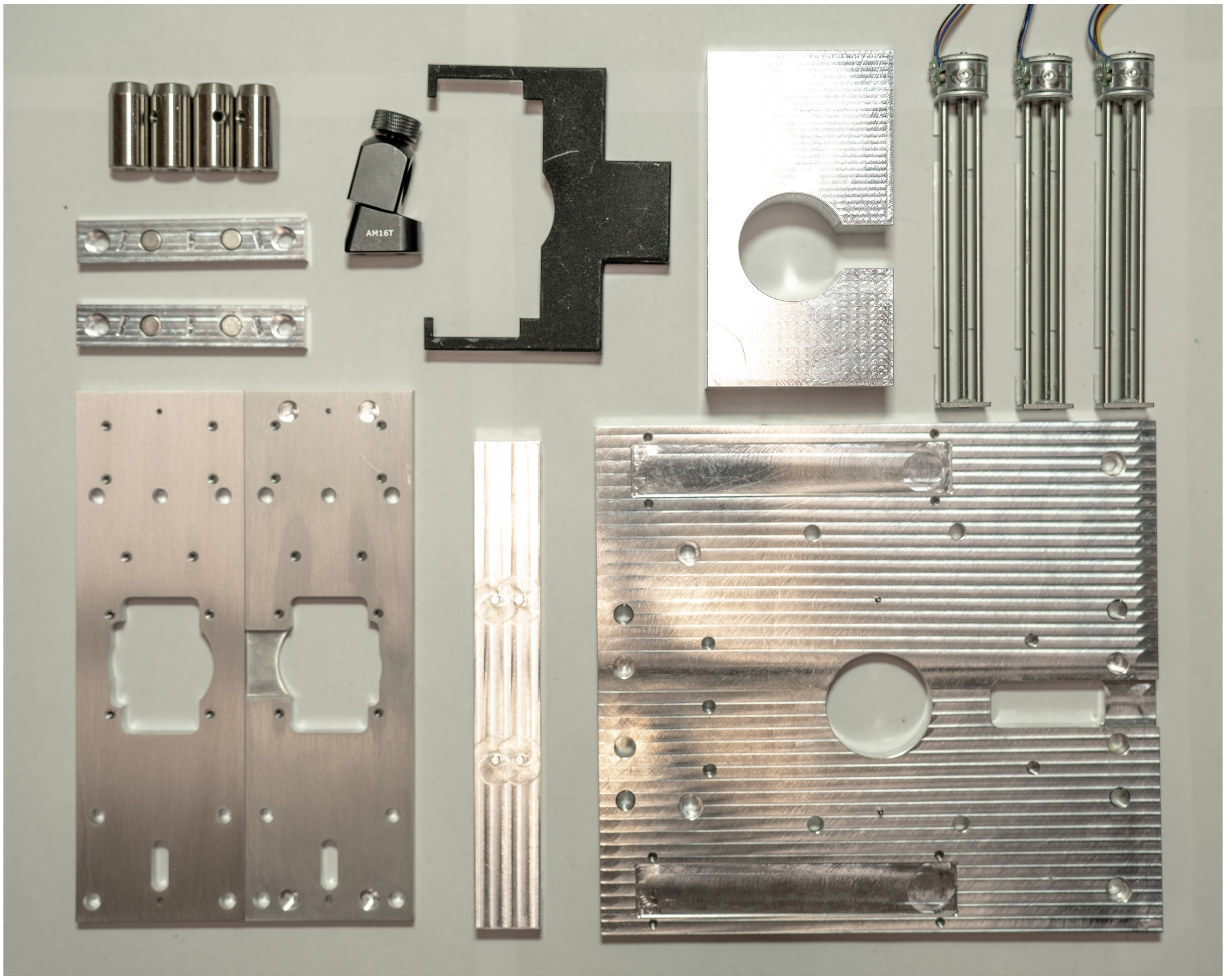

**Figure S9** Parts common to different configurations of *octopi* presented. These parts constitute the scanning module and backbone of different illumination and imaging modules.

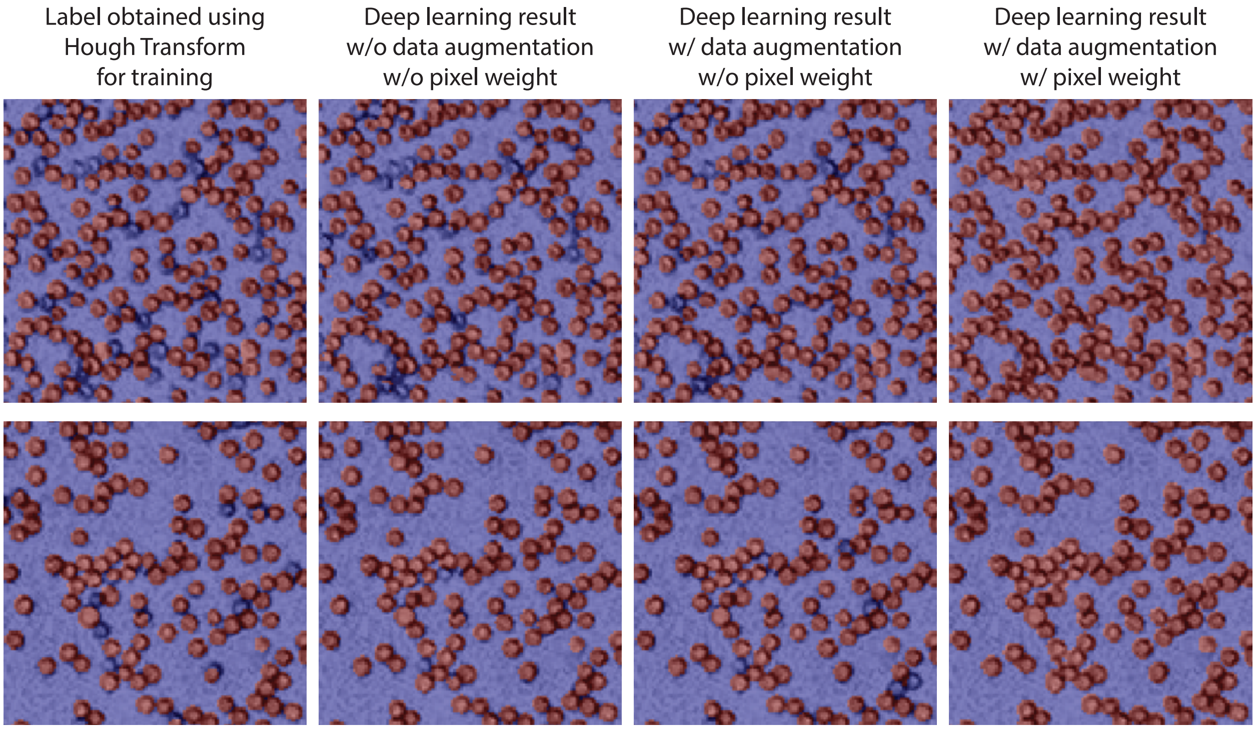

**Figure S10** Training of the convolutional neural network. As can be observed, data augmentation and introduction of class weights improves the performance of the neural network.

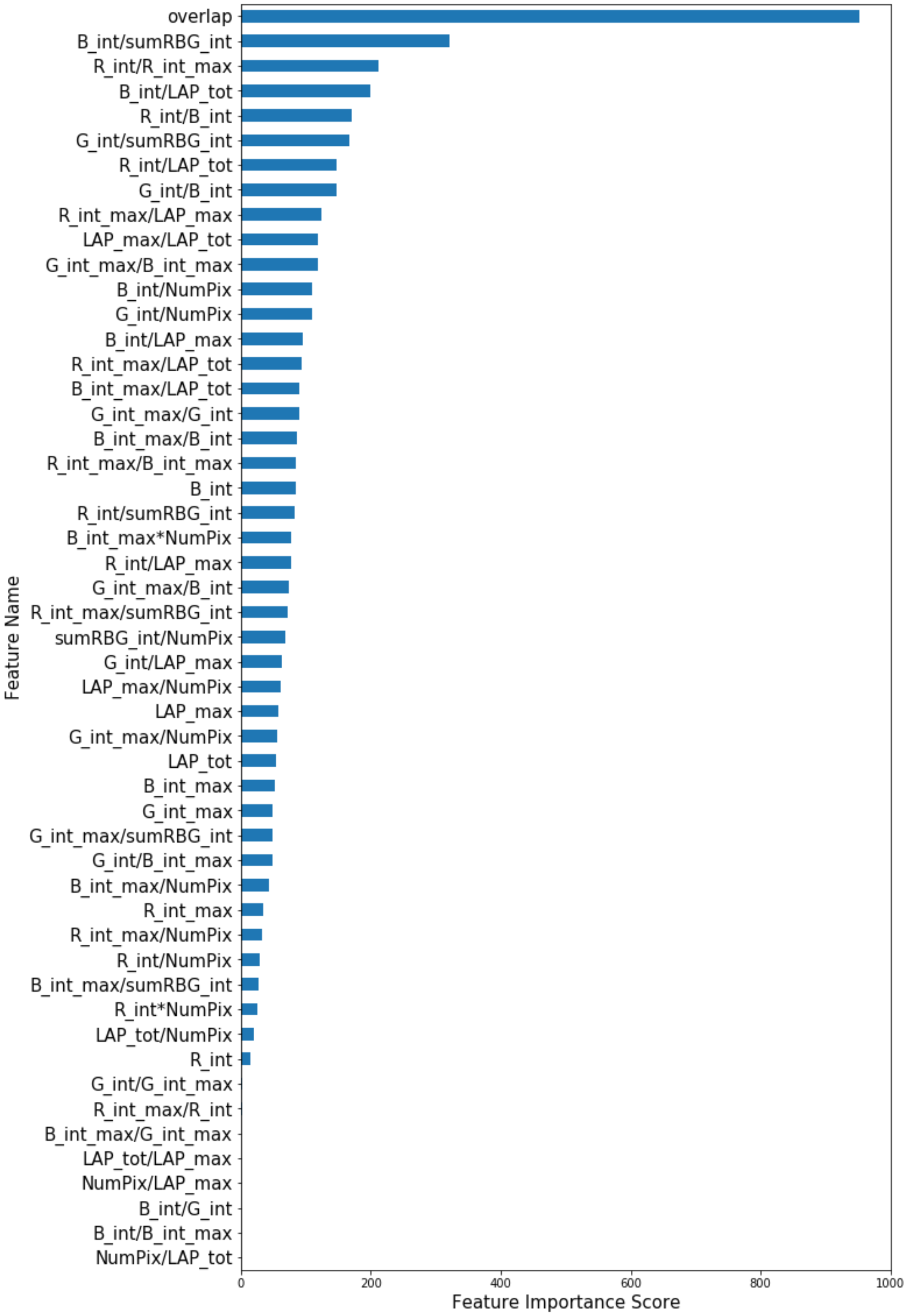

**Figure S11** Features used by the classifier and their importance scores output by the classifier in the training phase.

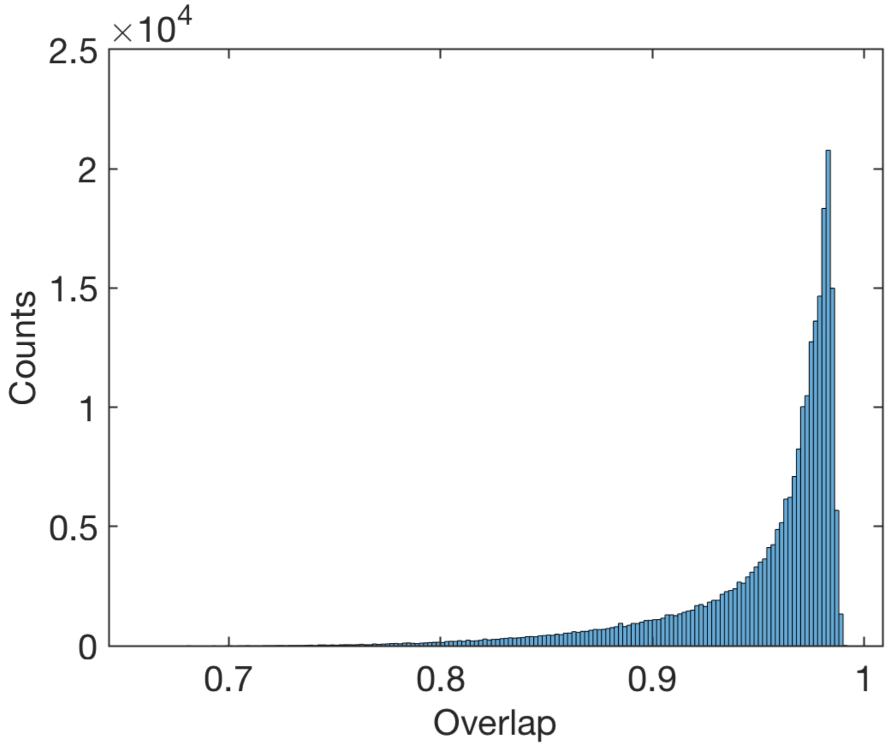

**Figure S12** Empirical distribution of overlap between red blood cells and *P. falciparum* parasites

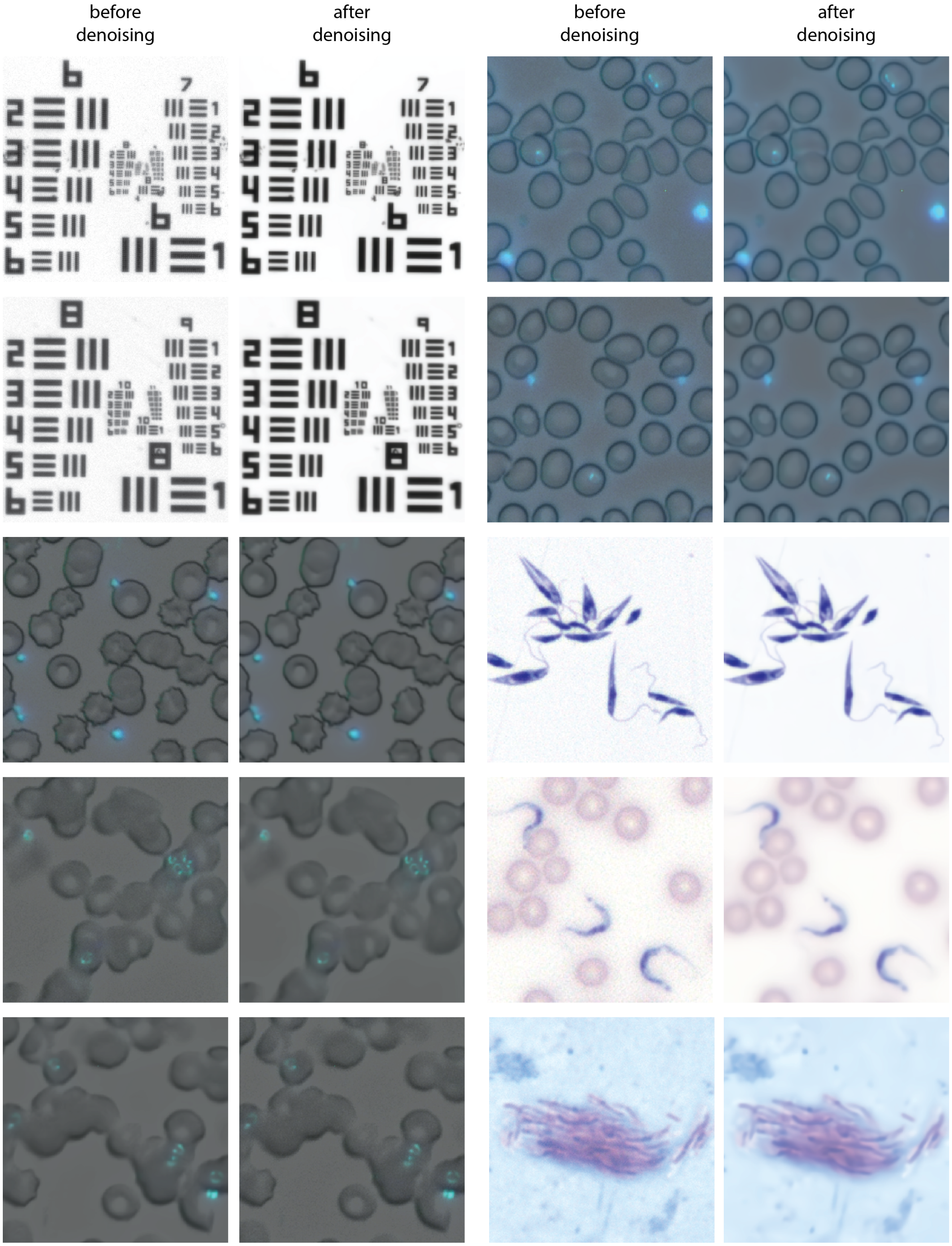

**Figure S13** Images before and after denoising. Denoising is done using FFDNet.

| **Module** | **Item** | **per unit cost @ 10 units** | **per unit cost @ 100 units** | **per unit cost @ 1000 units** |
|---|---|---|---|---|
| Scanner | Machined Parts | $180 | $85 | $55 |
| | Motors | $10 | $10 | $10 |
| | **Total** | $190 | $95 | $65 |
| Low mag imaging module | Machined parts | $130 | $54 | $34 |
| | Linear actuator | $100 | $60 | $31 |
| | Pi camera | $25 | $25 | $25 |
| | Cell phone lens | $4 | $4 | $4 |
| | Interference filter | $3 | $3 | $3 |
| | **Total** | $262 | $146 | $97 |
| Illumination module | Machined Parts | $80 | N/A | N/A |
| | LED panel | $2 | $2 | $2 |
| | Optics | $20 | $16 | $16 |
| | **Total** | $102 | $18 | $18 |
| Laser module | Laser | $7 | $5 | $5 |
| Electronics | Electronics | $25 | $20 | $15 |
| Computation module | Raspberry Pi | $35 | $35 | $35 |
| **Total Cost** | | **$621** | **$319** | **$235** |
| High mag imaging module | Machined parts | $120 | $60 | $40 |
| | Linear stage | $55 | $54 | $52 |
| | Piezo stacks | $62 | $44 | $35 |
| | Piezo driver | $60 | $15 | $10 |
| | Pi camera | $25 | $25 | $25 |
| | M12 lens | $5 | $5 | $5 |
| | Interference filter | $25 | $10 | $10 |
| | Objective (40x) | $60 | $50 | $50 |
| | **Total** | $412 | $263 | $227 |
| additional 100x objective | 100x objective | $73 | $60 | $60 |
| Computation module | Jetson Nano | $99 | $99 | $99 |

**Table. S1** Estimated cost breakdown for *Octopi* per number of units produced.
